# Multi-omics analyses and machine learning prediction of oviductal responses in the presence of gametes and embryos

**DOI:** 10.1101/2024.06.13.598905

**Authors:** Ryan M. Finnerty, Daniel J. Carulli, Akshata Hegde, Yanli Wang, Frimpong Baodu, Sarayut Winuthayanon, Jianlin Cheng, Wipawee Winuthayanon

**Author notes:** **Corresponding author**: Address: 1030 Hitt Street, Columbia, MO, 65211, USA. **Author Contributions:** R.M.F and W.W. designed the experiments. R.M.F. D.J.C., A.H., Y.W., F.B., and S.W. performed the experiments, A.H., Y.W., F.B., and J.C. designed machine learning methods. R.M.F., D.J.C., A.H., Y.W., F.B., S.W., J.C., and W.W. analyzed the data and wrote, edited, reviewed, and approved the final version of the manuscript. **Classification:** Major: Biological Sciences, Minor: Developmental Biology.

## Abstract

The oviduct is the site of fertilization and preimplantation embryo development in mammals. Evidence suggests that gametes alter oviductal gene expression. To delineate the adaptive interactions between the oviduct and gamete/embryo, we performed a multi-omics characterization of oviductal tissues utilizing bulk RNA-sequencing (RNA-seq), single-cell RNA-sequencing (scRNA-seq), and proteomics collected from distal and proximal at various stages after mating in mice. We observed robust region-specific transcriptional signatures. Specifically, the presence of sperm induces genes involved in pro-inflammatory responses in the proximal region at 0.5 days post-coitus (dpc). Genes involved in inflammatory responses were produced specifically by secretory epithelial cells in the oviduct. At 1.5 and 2.5 dpc, genes involved in pyruvate and glycolysis were enriched in the proximal region, potentially providing metabolic support for developing embryos. Abundant proteins in the oviductal fluid were differentially observed between naturally fertilized and superovulated samples. RNA-seq data were used to identify transcription factors predicted to influence protein abundance in the proteomic data via a novel machine learning model based on transformers of integrating transcriptomics and proteomics data. The transformers identified influential transcription factors and correlated predictive protein expressions in alignment with the *in vivo*-derived data. Lastly, we found some differences between inflammatory responses in sperm-exposed mouse oviducts compared to hydrosalpinx fallopian tubes from patients. In conclusion, our multi-omics characterization and subsequent *in vivo* confirmation of proteins/RNAs indicate that the oviduct is adaptive and responsive to the presence of sperm and embryos in a spatiotemporal manner.

**Significance Statement:** We conducted a detailed molecular study of how the oviduct changes its gene expression and protein production in response to sperm and embryos after mating in mice. We found that the oviduct has distinct molecular signatures in different regions – upper versus lower regions. Shortly after mating, inflammatory responses are turned on in the lower regions due to the presence of sperm. A day later, metabolic genes ramp up in the lower regions, likely to provide nutrients for the developing embryos. Overall, this multi-omics study revealed that the oviduct dynamically adapts its molecular makeup over time and space to accommodate and support sperm, eggs and embryos.

## INTRODUCTION

Optimal physiological conditions in the oviduct (fallopian tube in humans) provide an adaptive microenvironment for several reproductive processes ranging from sperm capacitation and transport to fertilization and embryonic development (1). The oviduct comprises four main regions: infundibulum (responsible for oocyte pick-up), ampulla (site of fertilization), isthmus (sperm capacitation/transport and preimplantation embryonic development), and the uterotubal junction (UTJ; responsible for filtering sperm and embryo transit to the uterus). Several studies demonstrated that distal (infundibulum and ampulla: IA) and proximal (isthmus and UTJ: IU) regions of the oviduct have distinct transcriptional profiles (2–5).

However, it is unclear how the presence of the sperm and embryo(s) modulates the oviductal responses. The presence of gametes and embryos has been shown to alter gene expression in secretory and ciliated cells of the oviduct during the preimplantation period (6–9). Additionally, it was reported that the endometrium responded differently to *in vivo-*derived embryos compared to embryos derived from *in vitro* fertilization (IVF) or somatic cell nuclear transfer in large animal models (10, 11), suggesting a maternal response to the presence of different types of embryos. Indeed, variations in the relative abundance of sets of genes involved in compaction and cavitation, desmosomal glycoproteins, metabolism, mRNA processing, stress, trophoblastic function, and growth and development have been observed in *in vitro*-produced embryos compared to their *in vivo* counterparts (12–15). Lastly, a growing consensus in several species indicates that epigenetic events in preimplantation embryos contribute to altered developmental potential both early and later in life (16).

Reciprocal embryo-oviduct interactions stem largely from investigating oviductal transport of fertilized/unfertilized embryos/oocytes in livestock (6, 7, 17–30) and rodents (2, 8, 31, 32). In humans, an embryo-derived platelet-activating factor has been implicated in the control of embryo transport to the uterus (33). It has been suggested that fertilized embryos produce prostaglandin E2 that facilitates transport to the uterus in mares (23, 25), whereas non-fertilized eggs remain in the oviduct (17). In hamsters, fertilized embryos are transported more expeditiously to the uterus compared to unfertilized eggs (31). In rats, transferred advanced-stage embryos (4-cell vs 1-cell) arrive in the uterus prematurely (32). In pigs (7) and cows (6), proinflammatory responses in the oviduct are down-regulated by the presence of embryos, suggesting that the embryo may facilitate maternal embryo tolerance during its passage through the oviduct. However, alterations in the oviductal transcriptome are difficult to detect in mono-ovulatory species (22, 25) indicating that the effect of the embryo in the oviduct is localized.

As for the sperm, observations suggest a filtering process as sperm migrates from the uterus, through the UTJ, into the oviduct (34, 35). After entering through the UTJ, sperm interact with ciliated cells in the isthmus to form a reservoir, undergo capacitation and are subsequently released to initiate the acrosomal reaction prior to reaching the ampulla (35–37). However, sperm are allogenic to the female reproductive tract, as sperm have been observed to induce pro– and anti-inflammatory responses in the oviduct (38, 39). Additionally, phagocytic bodies in the luminal fluid at the isthmus region can engulf sperm for degradation in mice (40). In addition to sperm selection, the oviduct seemingly provides beneficial chemical and mechanical mechanisms through rheotaxis, thermotaxis, and chemotaxis that assist sperm in transportation and fertilization (41–43). These observations suggest that the oviduct provides a malleable environment that is plastic and adaptable to select and facilitate the fittest sperm for fertilization. Therefore, our study also intends to provide a better understanding of the oviductal environment before, during, and after the presence of sperm in different regions of the oviduct.

In recent years, the field of reproductive biology has increasingly leveraged artificial intelligence (AI) and machine learning technologies to delve deeper into the intricate mechanisms governing fertilization and embryonic growth. AI predictive models, such as powerful transformer models, have shown remarkable capabilities in analyzing large-scale biological data, encompassing multi-omics data, to unveil patterns and forecast outcomes with elevated precision (44). One of the critical attributes of transformer models is the attention mechanism, which empowers the model to focus on pertinent essential segments of the input data that are critical for predicted outcomes (45). This functionality proves advantageous in the domain of reproductive biology, wherein complex interplays among genes, proteins, and other biomolecules dictating fertility outcomes may be revealed by the attention mechanism. The objective of this investigation is to amalgamate a multi-omics strategy with a transformer-based AI predictive model to elucidate the adaptive characteristics of the oviduct during natural fertilization.

Based on this premise, our study aims to elucidate the adaptive nature of the oviduct using a multi-omics approach during natural fertilization and preimplantation embryo development in a mouse model. We dissected oviducts from naturally fertilized mice at 0.5, 1.5, 2.5, and 3.5 days post-coitus, pseudopregnancy, and superovulation (dpc, dpp, SO, respectively). Gene expression profiles were analyzed from two different regions of the oviduct (IA and IU) using bulk-RNA and single-cell RNA (scRNA) sequencing (seq) analyses, generating a spatiotemporal depiction of gene expression in the oviduct.

Observations of RNA expression profiles from bulk RNA-seq findings were reinforced by scRNA-seq and LC-MS/MS proteomics analysis. Lastly, we integrated bulk RNA-seq and proteomics datasets to develop the initial stages of a machine-learning predictive model, which can identify influential transcription factors and correlate predictive protein expressions based on *in vivo*-derived data. Overall, we observed a robust transition of transcripts in the oviduct after sperm exposure at 0.5 dpc to other timepoints during preimplantation in both IA and IU regions. One of our key observations, was an elevated proinflammatory transcriptional and proteomic profile at 0.5 dpc, likely due to the presence of sperm preceding an anti-inflammatory condition 24 hrs later, correlating with the spatial presence of the embryo in the IU region at 1.5 dpc. Furthermore, this study paves the way for formulating a pioneering integrative AI model methodology tailored to integrate transcriptomics and proteomics data.

## RESULT

### Bulk RNA-seq analysis reveals a dynamic transcriptional profile during pregnancy that exhibits a distinct signature from pseudopregnancy

To ensure the presence and location of embryos/eggs in the oviduct in our model, we sampled the oviduct at different timepoints and evaluated the location of the embryos/ovulated eggs using H&E staining. Fertilized and unfertilized eggs with surrounding cumulus cells were in the ampulla at 0.5 dpc/dpp, respectively (Fig. 1*A*). Two-cell embryos and unfertilized eggs were clustered in the isthmus at 1.5 dpc/dpp. At 2.5 dpp/dpc, unfertilized eggs and embryos at the 8-cell to the morula stage were halted in a single-file formation at the UTJ region. At 3.5 dpc, the UTJ region was devoid of embryos/oocytes as all embryos/oocytes were transported to the uterus and, therefore, not included in the figure.

**Fig. 1.**
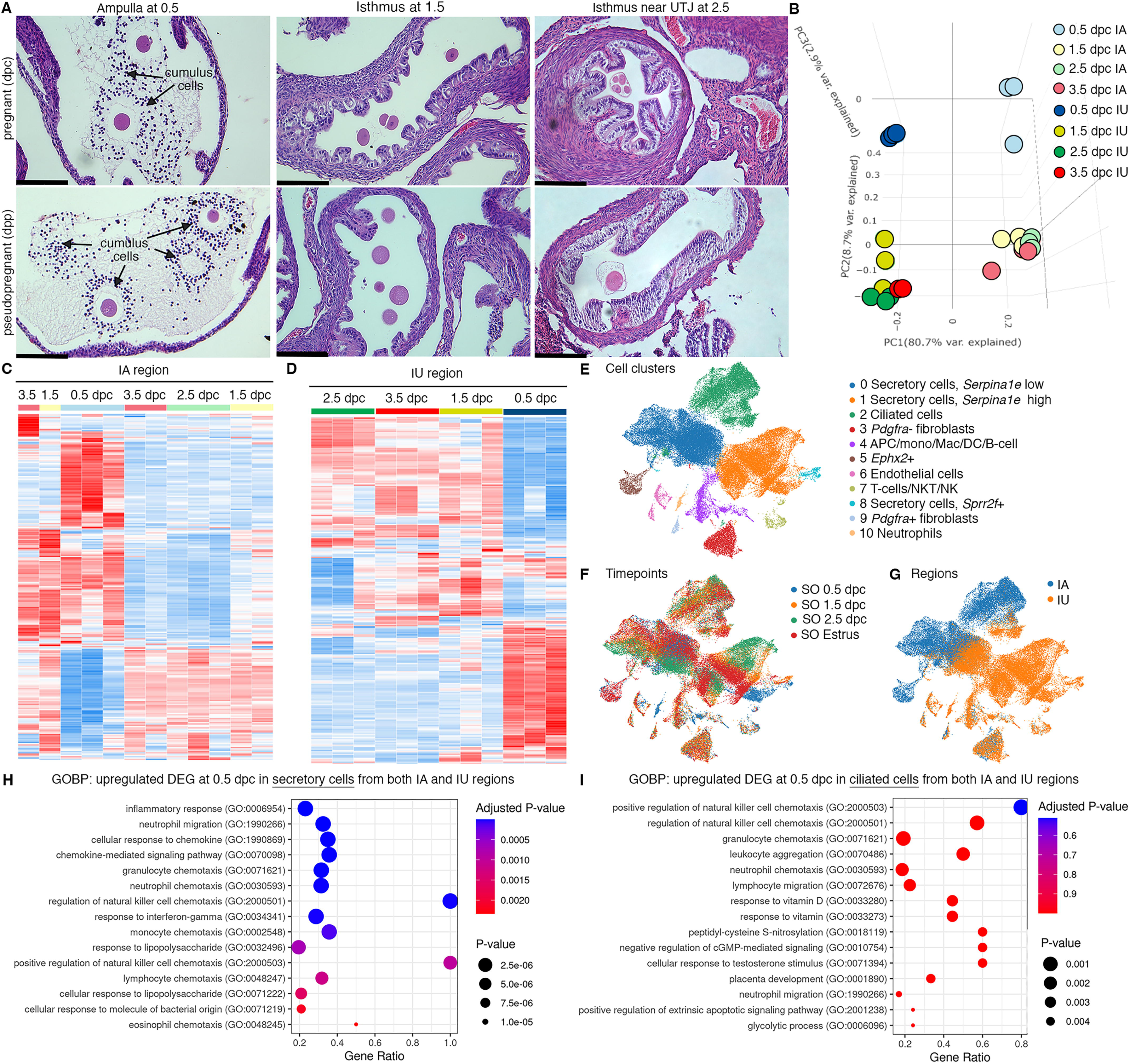
Transcriptomic analyses of the oviduct at different stages of early pregnancy. **A)** Histological analysis of different oviductal regions (ampulla, isthmus, and near the uterotubal junction (UTJ)) in mice at different stages of pregnancy (0.5, 1.5, and 2.5 dpc) and pseudopregnancy (0.5, 1.5, and 2.5 dpp) using H&E staining (scale bars = 132 μm, n=3 mice/timepoint/region). Arrows indicate cumulus cells surrounding the eggs/fertilized eggs called cumulus-oocyte complexes. **B)** Principal Component Analysis (PCA) of top 2500 DEGs identified from bulk-RNA seq of the infundibulum+ampulla (IA) and isthmus+UTJ (IU) regions of the oviduct collected at 0.5, 1.5, 2.5, and 3.5 dpc. Heatmap plots of unsupervised hierarchical clustering of top 2500 DEGs identified from bulk-RNA seq in the oviduct during pregnancy (0.5, 1.5, 2.5, and 3.5 dpc) of **C.** IA and **D.** IU regions. **E-F)** scRNA-seq analysis of the oviduct from superovulated (SO) estrus, SO 0.5 dpc, SO 1.5 dpc, and SO 2.5 dpc. Uniform Manifold Approximation and Projection (UMAP) of **E)** cell clusters identified from the oviduct **F)** at different regions (IA and IU) and **G)** at different timepoints (n=3-4 mice/timepoint/region). **H** and **I.** GOBPs dot plots of scRNA-seq analysis when compared between upregulated DEGs from **H.** secretory epithelial cells and **I.** ciliated epithelial cells at SO 0.5 dpc compared to SO Estrus from both IA and IU regions.

To determine whether the transcriptional profiles of each oviductal region are unique at fertilization and different developmental stages during preimplantation development, bulk RNA-seq analysis was performed at 0.5, 1.5, 2.5, and 3.5 dpc. Additionally, we aim to address whether changes in transcriptional signatures in the oviduct are governed by hormonal fluctuations or the presence of sperm/embryos/eggs. Therefore, oviducts from females at corresponding days post-mating with vasectomized males at (0.5, 1.5, 2.5, and 3.5 dpp) were used for comparisons. PCA plots were generated using the top 2,500 differentially expressed genes (DEGs, Fig. 1*B* and Fig. S1 *A* and *B*). Broad observations of region-specific transcriptome uniqueness exhibited segregation of all IA and IU biological replicates to opposite ends of the center axis on the PC1, reinforcing previous findings (5) that IA and IU regions behave differently with respect to transcriptional activity. Surprisingly, with respect to both the IA and IU regions, overall transcripts at 0.5 dpc (Fig. 1*B*) were segregated to the topmost axis along the PC2 plane, while 1.5-3.5 dpc biological replicates were segregated to the bottommost axis along the PC2 plane.

Expression signatures of the top 2500 DEGs in the IA region during pseudopregnancy were similar to those during pregnancy, as indicated by a heatmap generated using unsupervised hierarchical clustering (Fig. S1 *C* and *D*). However, there were exceptions at 0.5 (Fig. S1*C*, blue box) and 1.5 (Fig. S1*C*, black box) dpc/dpp. Unlike the IA region, DEGs in the IU region were more dynamic, as indicated by the presence of unique sets of genes at 0.5, 1.5, 2.5, and 3.5 dpc timepoints between pregnancy vs pseudopregnancy (Fig. 1*D* and Fig. S1*D*, blue, black, and red boxes). These findings indicate that oviduct transcripts from pregnant mice also possessed distinct signatures from pseudopregnant samples. Overall, data suggest that the transcriptional profile in the oviduct at all stages during the preimplantation period in the IU region is more dynamic compared to the IA region.

### Cellular responses to inflammation are enriched at the proximal (IU) and distal (IA) regions in response to the sperm

Oviductal transcription signatures were more unique at 0.5 dpc when compared to those at 1.5-3.5 dpc (i.e., 0.5 dpc vs. rest) in both IA and IU regions (Fig. 1 *C* and *D*). To determine the biological process of genes that were differentially expressed at 0.5 dpc compared to 1.5-3.5 dpc in both IA and IU regions, an initial analysis (0.5 dpc vs. rest) was chosen to isolate what distinct processes may be occurring during the transition from 0.5 dpc. Unique DEGs upregulated at 0.5 dpc in the IA region were enriched for the following biological processes (BPs): extracellular matrix (ECM) organization, extracellular structure organization, collagen fibril organization, and Ca^2+^ ion homeostasis, among others (Fig. S2*A*). Most interestingly, we found upregulated DEGs enriched for BPs at 0.5 dpc in the IU region included cellular response to cytokine stimulus, neutrophil migration, response to interferon-gamma, response to lipopolysaccharide, and neutrophil chemotaxis (Fig. S2*B*). Moreover, there are multiple BPs involved in the glucose catabolic process to pyruvate, in addition to other pyruvate metabolic processes that were uniquely upregulated at 0.5 dpc in the IU region (Fig. S2*B*). Next, we evaluated the IA region at 0.5 dpc compared to 1.5 dpc (presence of sperm vs. 24 h post-sperm exposure in the presence of embryos). We observed significant BPs that were enriched for downregulated DEGs at the IA region at 1.5 dpc compared to 0.5 dpc (Fig. S2 *C* and *D*). These processes included response to interferon-gamma, neutrophil chemotaxis, cytokine-mediated signaling pathway, and neutrophil migration. BPs common to the IA region at 0.5 dpc also included ECM organization, extracellular structure organization, and collagen fibril organization.

DEGs were more dynamic in the IU region during preimplantation embryo development compared to the IA region. At 0.5 dpc, the sperm are present, creating a sperm reservoir in the isthmus (46). When comparing 0.5 dpc to 0.5 dpp in the IU region (presence or absence of sperm, respectively), gene ontology biological processes (GOBP) analysis revealed significant enrichment of multiple proinflammatory BPs, including inflammatory response, neutrophil migration, neutrophil chemotaxis, regulation of phagocytosis, positive regulation of acute inflammatory response, and response to lipopolysaccharide when sperm were present in the IU (Fig. S2 *E* and *F*). Therefore, it is likely that, at 0.5 dpc, the isthmus region of the oviduct was heavily regulated for an inflammatory response in the presence of sperm while simultaneously preparing for the metabolic switch of the embryos from pyruvate to glucose metabolism.

Next KEGG analysis was used to determine molecular players; we found that genes in the tumor necrosis factor (TNF) signaling pathway were mostly upregulated at 0.5 dpc compared to 0.5 dpp at the IU region (Fig. S3*A*). Subsequently, an analysis comparing 0.5 dpc to 1.5 dpc in the IU region (presence of sperm vs. presence of embryos) demonstrated the most striking results. We found that most upregulated genes at 0.5 dpc were now downregulated at 1.5 dpc (Fig. S3*B*). Many of these genes are involved in the cellular response to cytokine stimulus, response to interferon-gamma, response to lipopolysaccharide, and neutrophil chemotaxis. These data strongly suggest that the oviduct may suppress the response to inflammation in the isthmus once the sperm is cleared to become conducive for the embryo’s survival at 1.5 dpc.

### scRNA-seq reveals that secretory epithelial cells contribute to the pro– and anti-inflammatory responses in the oviduct

To determine the key cell types responsible for the oviductal response to sperm/embryos, scRNA-seq analyses were leveraged. As we did not observe significant transcriptional changes from bulk RNA-seq at 3.5 dpc and embryos were not present in the oviduct at 3.5 dpc, we opted not to assess a 3.5 dpc timepoint in our scRNA-seq analysis. In this experiment, superovulation (SO) using exogenous gonadotropins was used due to technical limitations of sample collection for single-cell processing. Non-mated SO estrus samples were used as controls. First, we confirmed that all cell types previously reported (3) were present in the oviduct (Fig. 1*E* and *F*). There was minimum overlap between cells isolated from IA or IU regions (Fig. 1*G*). In addition, all cell types were present at all timepoints except for an *Ephx2*+ cluster (only present at SO 0.5 dpc and SO estrus) and a neutrophil cluster (*Ly6g+,* only present at SO 0.5 dpc).

Next, we investigated whether our findings from bulk RNA-seq data would be recapitulated in the scRNA-seq dataset. Here, we exclusively focused on the IU region as it was the most dynamically regulated region during early pregnancy. Based on GO BP analysis from bulk RNA-seq findings, we further assessed several genes that were upregulated at 0.5 and 1.5 dpc corresponding to GOBP terms ‘inactivation of mitogen-activated protein kinase (MAPK) activity’ and ‘MAP kinase phosphatase activity’. Genes associated with these pathways were mostly upregulated at SO 0.5 and SO 1.5 dpc (Fig. S3*C*, green and teal bars) and downregulated in SO estrus and SO 2.5 dpc in the IU regions (Fig. S3*D*, red and purple bars). These genes include dual-specificity phosphatase family (*Dusp1*, *Dusp5*, *Dusp6*, *Dusp10*), *Fos*, interleukin 1b (*Il1b*), IL1 receptor 2 *(Il1rb*), and others. As DUSP proteins are crucial for controlling inflammation and antimicrobial immune responses (36), we performed qPCR analysis to confirm both our bulk RNA-seq and scRNA-seq data with respect to MAPK signaling pathways. We found that *Dusp5* was expressed at a significantly higher level at 0.5 dpc compared to 0.5 dpp while *Mapk14* (*p38a*) was significantly upregulated at 1.5 dpc compared to 0.5 dpc (Fig. S3 *E* and *F*). We also further assessed several genes from the GOBP term ‘neutrophil-mediated immunity’ to explain the appearance at SO 0.5 dpc and subsequent disappearance of the neutrophil cluster at SO 1.5 dpc, respectively (Fig. 1*E* and *F*). Interestingly, these genes were found to be downregulated in both the IU and IA regions at the 1.5 and 2.5 dpc timepoints (Fig. S3 *G* and *H*, teal and purple bars).

To identify which cell population is contributing to the observed pro– and anti-inflammatory response in both IA and IU regions at SO 0.5 dpc. We evaluated DEGs in each cell population and performed GOBP analysis. Upregulated DEGs from secretory epithelial cells (both clusters 0 and 1; Fig 1 *E*-*I*) from both IA and IU regions at 0.5 dpc were enriched for BPs involved in inflammatory response, neutrophil migration, cellular response to chemokine, chemokine-mediated signaling pathways, and several others chemokine signaling pathways (Fig 1*H*). In contrast, when evaluated for upregulated DEGs in ciliated epithelial cells, similarly enriched biological processes were present, albeit in a less significant manner (Fig 1*I*). Therefore, it suggests that secretory cells are the key modulators responsible for the regulation of pro– and anti-inflammatory responses during pregnancy establishment.

### Oviductal luminal proteomics are dynamic at different preimplantation stages and SO exacerbates the transcriptional profile at each timepoint

To validate our transcriptomics data at a translational level, LC-MS/MS proteomic analysis was performed on secreted proteins in the oviductal luminal fluid at estrus, 0.5, 1.5, and 2.5 dpc. Note that proteomic analysis was not performed at 3.5 dpc as the embryos have vacated the oviduct at this stage. Additionally, we aim to address whether changes in proteomic profiles in the oviduct are governed by hormonal fluctuations. Oviductal luminal fluid was also collected at different stages after superovulation, including SO estrus, SO 0.5, SO 1.5, and SO 2.5 dpc. In agreement with the transcriptomic data, secreted proteins from 0.5 dpc and SO 0.5 dpc were segregated from all other timepoints (Fig. 2 *A*-*C*). Another difference was observed between 1.5 dpc and SO 1.5 dpc, at which 1.5 dpc proteomic dynamics correlated more with estrus and SO estrus biological replicates, while SO 1.5 dpc correlated more with 2.5 dpc and SO 2.5 dpc.

**Fig. 2.**
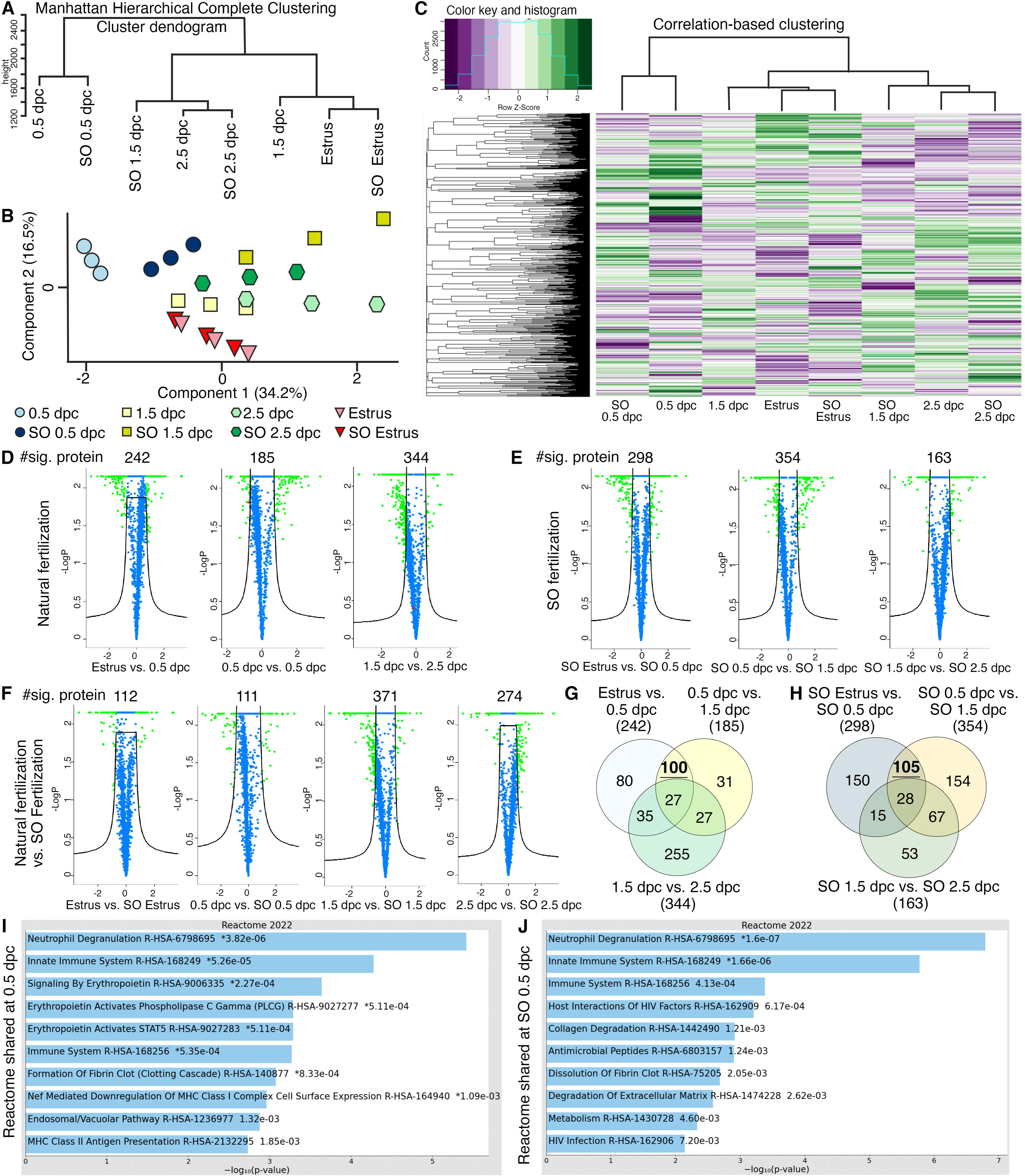
Analyses of protein abundance in the oviduct luminal fluid at different stages of pregnancy compared to Estrus. **A)** Manhattan hierarchical complete clustering dendrogram of natural (Estrus, 0.5 dpc, 1.5 dpc, and 2.5 dpc) and superovulated (SO Estrus, SO 0.5 dpc, SO 1.5 dpc, and SO 2.5 dpc) datasets (n= pooled of 3 biological samples/timepoint). **B)** PCA plot of all datasets generated utilizing Perseus software after integration of the Gaussian transformation. **C)** Correlation-based hierarchal clustering of all protein abundance. **D-F)** Volcano plots of significantly different protein abundances when compared between **D)** Natural fertilization, **E)** SO fertilization, and **F)** Natural fertilization vs. SO fertilization. Numbers of significant proteins were listed above the volcano plots. **G** and **H)** Gaussian transformed Perseus two-tail *t*-tests of differentially abundant proteins in oviductal fluid at different stages during **G)** Natural fertilization and **H)** SO fertilization. Differentially abundant proteins shared between Estrus and 0.5 dpc (100) or SO Estrus and SO 0.5 dpc (105) were underlined. **I** and **J**) Enricher Reactome pathway analysis of differentially abundant proteins shared at **I)** 0.5 and **H)** SO 0.5 dpc.

Analysis comparing naturally fertilized (dpc) pregnant samples yielded 242 differentially abundant proteins between Estrus and 0.5 dpc, 185 between 0.5 dpc and 1.5 dpc, and 344 between 1.5 and 2.5 dpc (Fig. 2*D*). Next, we elucidated whether SO treatment impacts protein secretion in the oviduct. There were 298, 354, and 163 differentially abundant proteins when compared between SO estrus vs. SO 0.5 dpc, SO 0.5 dpc vs. SO 1.5 dpc, and SO 1.5 dpc vs. SO 2.5 dpc, respectively (Fig. 2*E*). In addition, protein samples from naturally fertilized and SO samples were evaluated. There were 112 differentially abundant proteins between estrus and SO estrus, 111 between 0.5 dpc and SO 0.5 dpc, 371 between 1.5 dpc and SO 1.5 dpc, and 274 between 2.5 dpc and SO 2.5 dpc (Fig. 2*F*). These results indicate that luminal proteomics from the oviduct are dynamic during preimplantation development and SO stimulates the production and secretion of more abundant and unique proteins compared to the natural setting.

Next, we explored differentially abundant proteins commonly shared between estrus vs. 0.5 dpc and 0.5 dpc vs. 1.5 dpc. We found that a subset of shared 100 proteins were enriched for multiple pro-inflammatory Reactomes including neutrophil degranulation, innate immune system, and innate immune system (Fig. 2 *G* and *I*). In addition, when evaluated a subset of shared 105 protein samples with SO at the same timepoints, similar if not identical Reactomes occurred, with lower *p-*values (Fig. 2 *H* and *J*), indicating greater pathway enrichment in SO treatments. Lastly, differential protein abundance at 1.5 dpc and 2.5 dpc indicated the enrichment for Ras Homolog (RHO) GTPase signaling pathway and changes in epithelial remodeling (keratinization) (Fig. S4 *A* and *B*), respectively. Therefore, the pro-inflammatory Reactome profile appeared to have completely subsided at 2.5 dpc. These results reinforce our bulk and scRNA-seq observations of a pro-inflammatory condition occurring at 0.5 dpc. Moreover, SO conditions appear to exacerbate both expression abundance and the expression of additional unique proteins with respect to proinflammation when compared to naturally fertilized replicates.

### *In vivo* confirmation of identified multi-omics proinflammatory condition in the oviduct at 0.5 dpc

To validate the findings from multi-omics studies, we used RNAScope *in situ* hybridization staining of *Tlr2* (epithelium, stroma, and myosalpinx)*, Ly6g* (leukocytes), and *Ptprc* (common immune cell marker). We found a significant induction of *Tlr2* at 0.5 dpc compared to 0.5 dpp at the isthmus and UTJ regions (Fig. 3 *A* and *B*). *Ptprc+* and *Ly6g+* signals aggregated with greater intensities in the mesosalpinx, stromal layer, and blood vessels in the oviduct. Additionally, no positive *Ly6g+* cell expression was found in the luminal space of the oviduct, but rather restricted to stromal and epithelial cell linings along with blood vessels.

**Fig. 3.**
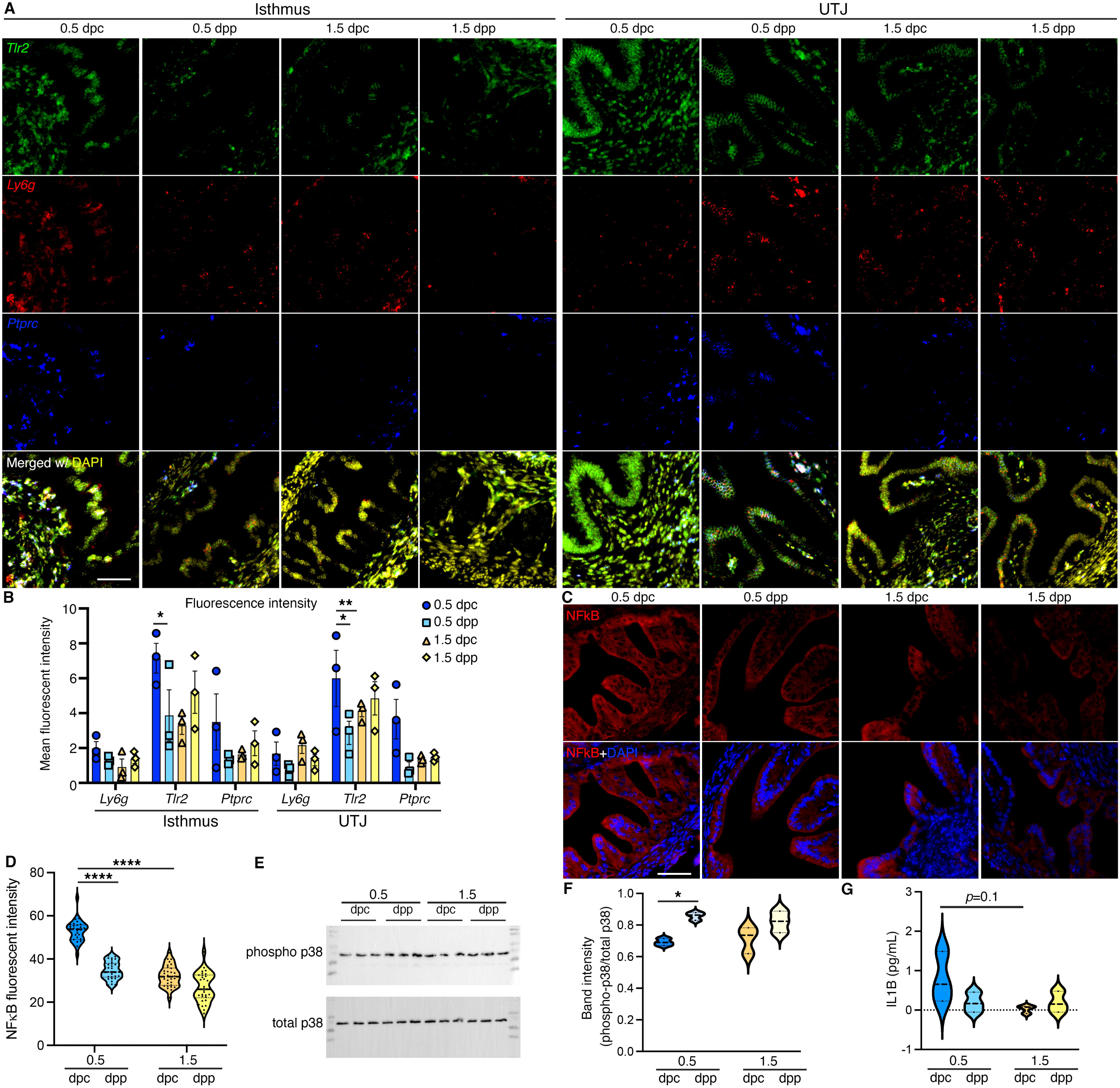
*In vivo* validation of RNA and proteins identified from bulk RNA– and scRNA-seq analysis. **A** and **B)** Expression of *Tlr2*, *Ly6g*, and *Ptprc* in the isthmus and UTJ regions at 0.5 dpc, 1.5 dpc, 0.5 dpp, and 1.5 dpp. Scale bar = 50 μm for all images in the panel. **B)** Quantification of fluorescent signal from images in **A** using FIJI software. Graph represent mean±SEM, n=3 mice/timepoint/region. **C)** Immunofluorescent staining of NFκB in the isthmus regions of the oviducts at 0.5 dpc, 1.5 dpc, 0.5 dpp, and 1.5 dpp. Scale bar = 50 μm for all images in the panel. **D)** Quantification of fluorescent signal from images in **C** using FIJI software. Violin plots represent all measurements, n=3 mice/timepoint/region, \*\*\*\**p*<0.001 compared to 0.5 dpc, unpaired *t-*test. **E)** Immunoblotting of phosphorylated p38 and total p38 in the whole oviduct collected at 0.5 dpc, 0.5 dpp, 1.5 dpc, and 1.5 dpp. **F)** Violin plots of the quantification of band intensity represented as phosphor-p38/total p38 ratio (n=3 mice/timepoint/region). \**p*<0.05 compared to 0.5 dpc, unpaired *t-*test. **G)** IL1β ELISA of protein from the whole oviduct at 0.5 dpc, 1.5 dpc, 0.5 dpp, and 1.5 dpp (n=3 mice/timepoint/region).

NFκB immunofluorescent staining was performed to evaluate the degree of inflammatory activation. The presence of NFκB appeared to be largely concentrated in the cytoplasm of all epithelial cells at all timepoints in the isthmus. The relative fluorescent signal was significantly greater at 0.5 dpc compared to 1.5 dpc or 0.5 dpp (Fig. 3 *C* and *D*). As p38 is the key mediator of the inflammatory response (47), we found that p38 and phosphorylated (p)-p38 proteins were expressed at all timepoints between 0.5 and 1.5 dpc and dpp (Fig. 3 *E* and *F*). Specifically, p-p38:total p38 ratio was significantly increased at 0.5 dpp compared to 0.5 dpc, suggesting an overall inflammatory response induced by mating regardless of the sperm exposure. In addition, the presence of pro-inflammatory cytokine, IL1β was evaluated. However, there was no difference in IL1β levels between timepoints (Fig. 3*G*). To summarize, these data suggest that an innate immune response occurs at 0.5 dpc in the isthmus and UTJ regions and that some of these responses were induced by the presence of seminal plasma regardless of the sperm.

### Integrating transcriptomics and proteomics data and identifying influential transcription factors in the oviduct via a predictive transformer model

Our machine learning method based on a transformer encoder model is rigorously evaluated against gene and protein expression data from 2.5 dpc of naturally fertilized samples, which was not used by the model during its training. The integrative transformer model was effective in predicting the protein abundance levels from bulk RNA-seq expression data with high accuracy. The evaluation results of the model are shown in Supplementary Table S1. The attention matrix for all genes against all proteins is extracted from the transformer model, which represents each gene’s potential influence level on the proteins (Fig. 4*A*). To focus on analyzing differentially expressed genes and proteins rather than all the genes and all proteins, differential gene expression and protein abundance expression between bulk RNA-seq and proteomic datasets at 0.5, 1.5, and 2.5 dpc were compared to Estrus and proteomics Estrus, respectively, followed by extraction of common significantly differentiated protein-coding genes or proteins (Fig. 4*B*). The differential gene expression is performed using DESeq2 (48) and the differential protein abundance analysis using Protrank (49). The top 25 “influential” transcripts (ITs) with the highest attention scores from all the transcription factors present in bulk RNA-seq data were extracted for every potentially influenced protein (IP) in the empirical proteomics datasets (Supplemental Datasets S1-S4).

**Fig. 4.**
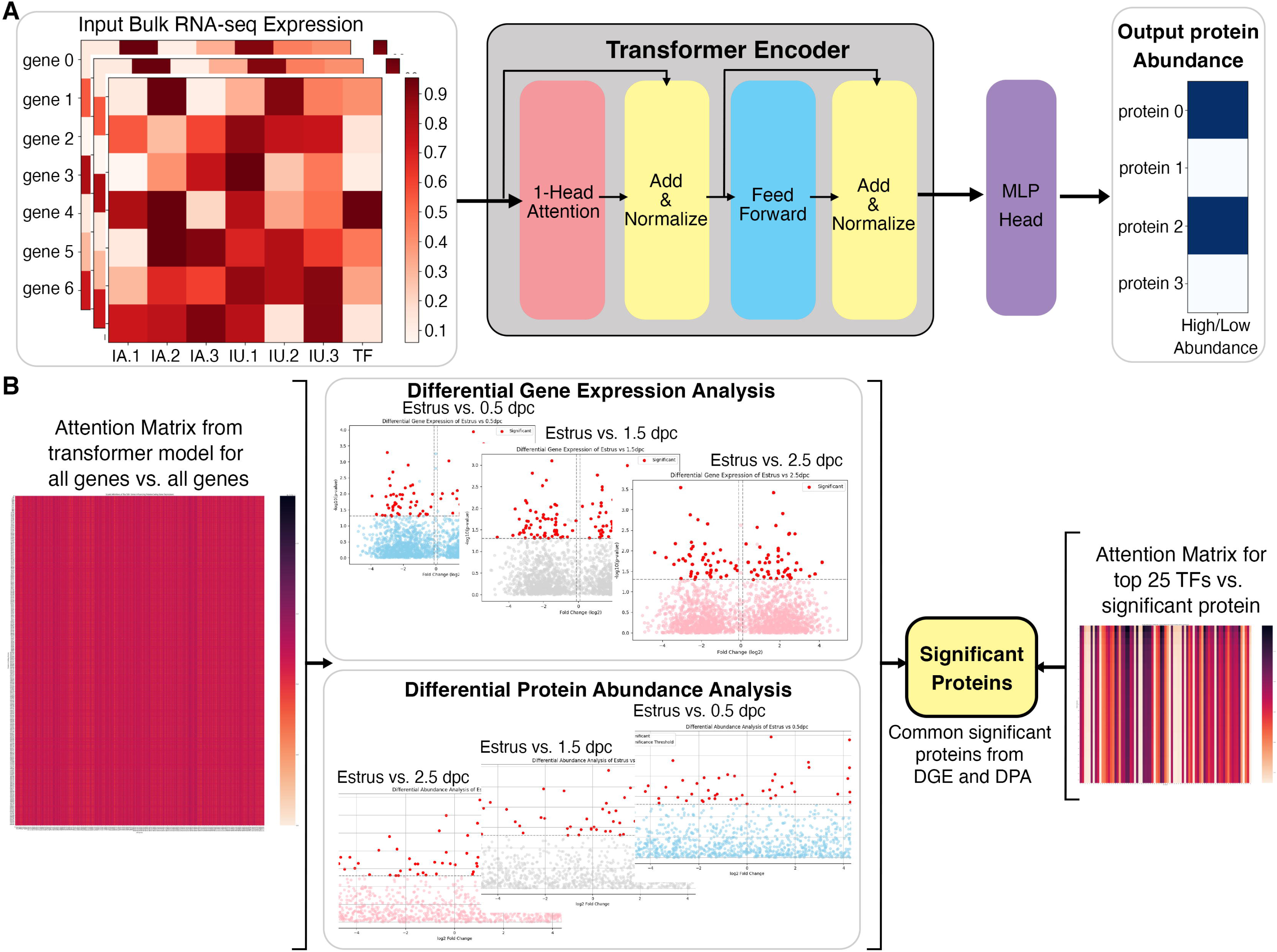
Overall architecture of the transformer-based model to predict proteomic abundance from bulk RNA-seq data of natural fertilization of oviduct. **A)** Preprocessing steps using bulk RNA-seq count per million (cpm) normalization to calculate expression values. The transformer model is equipped with a single-layer transformer encoder featuring a single-head (1-Head) self-attention mechanism to predict the abundancy of proteins (abundant or not) from the input RNA-seq data. “Head” refers to blocks, modules, or connections that perform specific tasks in neural networks. A specific threshold of 0.6/0.8 was defined to label proteins as high abundance or low abundance. The Multi-Layer Perceptron (MLP) Head refers to the output layer, which is designed to perform a classification task. In this model, The MLP layer uses a multi-layer perceptron or linear layer as the backbone to divide high abundance and low abundance based on the importance or attention weights given by the previous transformer layer. **B)** The visual representation of a method to extract the top 25 TFs for differential significant proteins. DGE; Differential gene expression, DPA; Differential protein abundance; TF, Transcription factors

The identified IT and IP lists were subsequently analyzed with Enrichr Reactome (2022) and GO Biological Process (2023) tools. A combination of both IT and IP lists generated function ontologies that match *in vivo* empirical observations. At 0.5 dpc, ITs predicted to influence protein abundance included, but are not limited to, *Clu, Anxa2, Nod2, Hspa8, Il17c, Il36b,* and *Il1b*, among many others (Supplemental Dataset S2). Among the top 25 ITs identified in high abundance at 1.5 dpc included *Cep126, Cfap126, Cfap54, Cfap65, Ift88, Ccdc40, Crocc2,* and *Clu* (Supplemental Dataset S3). Lastly, ITs abundant at 2.5 dpc included, but were not limited to, *Mapk15, Hsph1, Drc7, Togaram2, Tspan15, Igfbp2, Rnf112,* and *Traf3ip3* (Supplemental Dataset S4). Taken together, we have developed a predictive transformer model that has recapitulated a similar progressive observation as our *in vivo* empirically biological multi-omics model. Moreover, the predictive model suggests that ITs and IPs present at 0.5 dpc indicate a pro-inflammatory condition, followed by a shift to ciliogenesis and cellular stress maintenance at 1.5 dpc, subsequently initiating cellular homeostasis at 2.5 dpc. In addition, this predictive tool can be adapted to other biological disciplines to identify influential ITs and IPs using existing bulk RNA-seq databases. Overall, our study lays the groundwork for developing a novel and comprehensive AI model approach specifically designed to combine and predict influential ITs and IPs in biological samples.

### Evaluation of human hydrosalpinx Fallopian tubes compared to sperm-induced inflammation genes

To determine whether sperm-induced inflammatory responses in the mouse oviduct are similar to or different from human inflammation conditions, we reanalyzed publicly available scRNA-seq data from hydrosalpinx samples by Ulrich *et al* (50). We found that some of the sperm-induced inflammatory genes identified from our mouse study were present and upregulated in hydrosalpinx samples compared to healthy subjects (Fig. 5*A*). However, the differentially expressed levels, for example the *CCL2* gene, appeared to be marginal between healthy vs. hydrosalpinx samples (Fig. 5*B-C* and Supplemental Datasets S5). Nevertheless, the top five most enriched GOBPs related to inflammatory responses were Regulation of Complement Activation, Positive Regulation of Macrophage Migration Inhibitory Factor Signaling Pathway, MHC Class II Protein Complex Assembly, Positive Regulation of NK Cell Chemotaxis, and Negative Regulation of Metallopeptidase Activity (Fig. 5*D*). These GOBPs differed from those identified in mouse oviducts at 0.5 dpc, which were exposed to sperm enriched for neutrophil-related pathways, not macrophages or NK cells in hydrosalpinx samples.

**Fig. 5.**
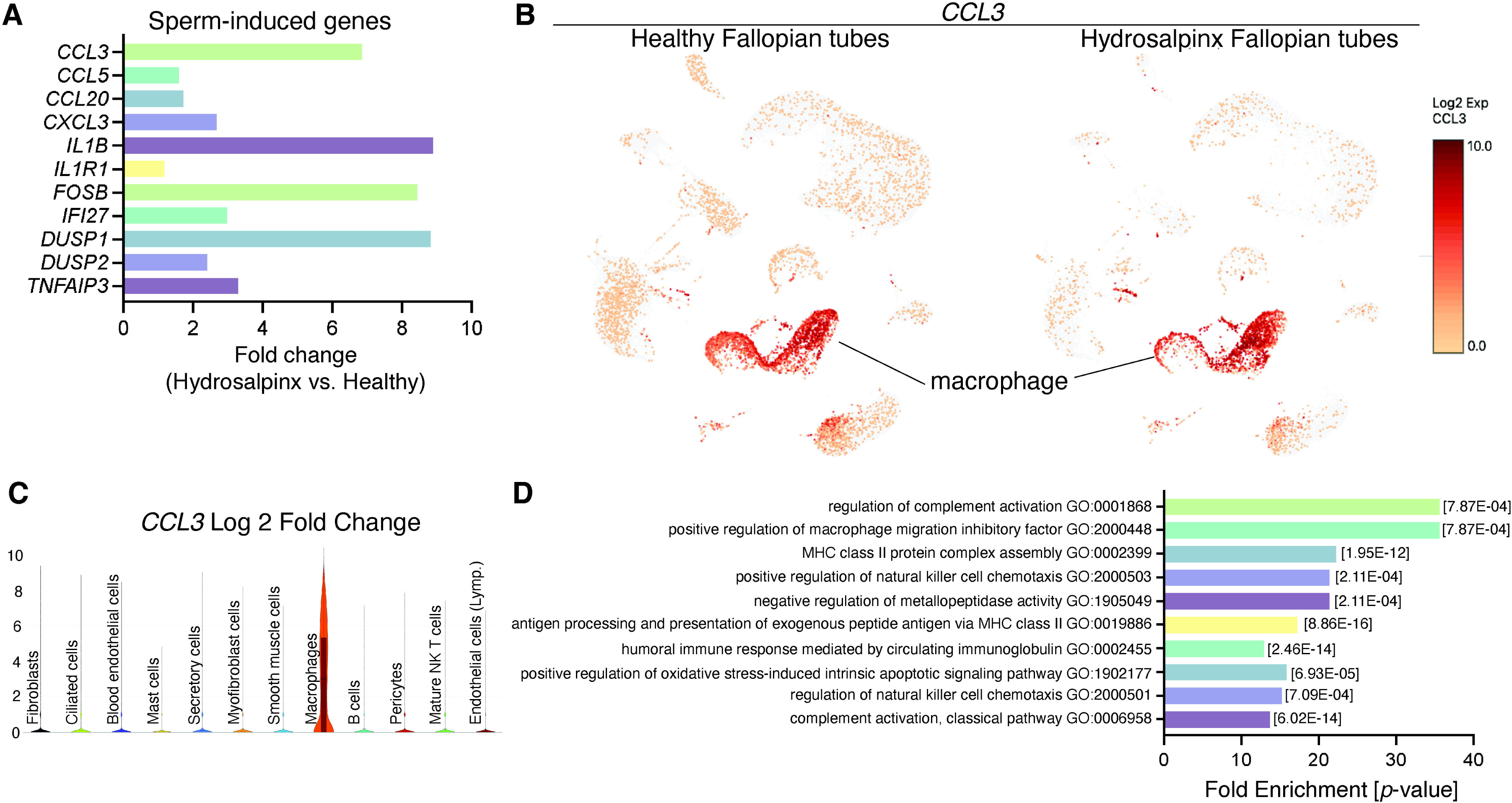
Reanalysis of data from Ulrich *et al.* using hydrosalpinx vs. healthy Fallopian tube samples from GSE178101. **A)** Expression of sperm-induced genes identified from this current study in the hydrosalpinx compared to healthy Fallopian tube samples. **B)** UMAP of *CCL3* in healthy and hydrosalpinx Fallopian tubes in the macrophage populations. **C)** Log2 Fold change of *CCL3* in a violin plot comparing hydrosalpinx vs. healthy Fallopian tubes. **D)** Enriched GOBPs related to inflammatory responses in the hydrosalpinx samples.

## DISCUSSION

Here, we performed the *in vivo* multi-omics characterization of the oviduct in the mouse model. We integrated a total of 68 biological samples using bulk-RNA sequencing (24 total biological replicates), scRNA-sequencing (20 total biological replicates), and LC-MS/MS (24 total biological replicates) analyses. In addition, our Bulk-RNA seq and proteomic data are immediately available to the scientific community in a web search format (details in Methods). Here, we reinforced significantly enriched pathways shared between different multi-omics techniques. We validated previous findings (5) that the transcriptional profile of the oviduct between the IA and IU regions is unique and region-specific based on PCA analyses. Both the IA and IU regions are most distinct at 0.5 dpc compared to other timepoints based on PCA analyses, with unique transcription patterns becoming most disrupted at 0.5 dpc in both pregnancy and pseudopregnancy datasets. Large sets of DEGs display a dramatic shift from either being up-or downregulated between 0.5 and 1.5 dpc in all –omics characterizations. The changes at 0.5 dpc appear to subside at 1.5-3.5 dpc, with fewer unique clusters of genes that were dynamic between timepoints, indicating that either the absence of sperm or the presence of embryos drives the oviduct transition.

However, the number of dynamic clusters of genes was greater in the IU region than in the IA region. This finding suggests that the IU region is more dynamic and responsive to the presence of gametes (sperm and oocytes/embryos) compared to the IA region.

At 0.5 dpc, we found that there were unique upregulated DEGs that corresponded with BPs involved in tissue remodeling and muscle filament sliding, such as ECM and collagen fibril organization. Wang and Larina showed that during this timepoint, ciliated epithelial cells in the ampulla region are responsible for creating a circular motion of the cumulus-oocyte complexes (COCs) within the ampulla (51). Here, using scRNA-seq analysis, we found that the ciliated cell population showed suppressed expression of genes involved in cilia assembly at SO 0.5 dpc and in SO Estrus (COCs are present in these two groups) in the IA region. This finding suggests that when the COCs are present in the ampulla, ciliated cells are functionally active. As SO results in higher levels of E_2_ due to increased mature follicles, ovulated eggs, and higher volume of follicular fluid, it is also likely that these changes after SO could lead to biological alterations observed in our study. Interestingly, the *Ephx2*+ cluster is mainly present in the SO 0.5 dpc and SO estrus samples. *Ephx2* encodes epoxide hydrolase 2, which converts epoxides to dihydrodiols. Recent findings suggest that EPHX2 may play a role in primary hypertension in humans (52). However, the reproductive-related functions of EPHX2 have not yet been investigated. Therefore, we believe this presents an opportunity for future research to define its role in preimplantation development as a result of SO.

The presence of sperm at 0.5 dpc strongly perturbed the IU region at 0.5 dpc, most likely due to a greater population of sperm in the IU, as a sperm reservoir (46, 53), compared to the IA region. This perturbation was minimally detected at 0.5 dpp. Multi-omics analysis and observations in bulk RNA-seq, scRNA-seq, and luminal proteomics datasets are in agreement with the previous finding (54) that seminal fluid and sperm may be the dominant influencers for stimulating inflammatory responsive pathways in the oviduct at 0.5 dpc. We also established here, for the first time, that these observed inflammatory responses may be facilitated by the secretory cell population in the IU region when compared to other cell types. DUSP proteins modulate inflammation and antimicrobial immune responses (36), and MAPK signaling pathways are involved in both pro– and anti-inflammatory pathways (36, 55). Therefore, we hypothesized that the observed inflammatory response was facilitated in part by the activation of MAPK signaling pathways, as indicated by a significant increase in expression of *Dusp5*, which was unique to the IU region after sperm exposure. Overall, the presence of sperm at 0.5 dpc induces a strong pro-inflammatory response in the IU region with upregulation of genes involved in inflammatory cytokines, neutrophil activation, lymphocyte recruitment and T-cell proliferation. ScRNA-seq data suggests that the oviduct is immunodynamic as indicated by the presence of immune cells as indicated by several immune markers such as neutrophils (*Ly6g*+), leukocyte (*Ptrpc*+), T cells (*Cd3d+, Cd3g+*), NK cells (*Nkg7+, Klrb1c+*), among others. This finding agrees with previous studies in human Fallopian tubes, as well as from our and other laboratories (3, 56, 57).

Sperm migration from the uterus through the UTJ into the oviduct has been an observable phenomenon dating back five decades (34). Additionally, phagocytic bodies engulfing sperm in mice luminal fluid in the isthmus region have also been observed in literature pre-dating the 1980s (40). The prevailing theory is that “fit” sperm display inherently, via intracellular processes and genetic cargos, membrane “passport” proteins that allow them to not only gain access through the UTJ, but also subsequently bind to the epithelium in the isthmus region (58). The *in situ* hybridization analysis of the IU region reinforces these observations, suggesting additionally that not only do sperm require specific membrane proteins to function properly in the uterus and oviduct, but also that sperm must evade phagocytosis from an innate immune response. Our findings showed that *Ptrprc*+ cells were present in the stromal and epithelial layers in the presence of sperm at 0.5 dpc in the UTJ. Similarly, a significant increase in *Tlr2+* cells was observed at the epithelial lining of the isthmus and UTJ regions. *Tlr2* is a part of the Toll-like receptor superfamily of proteins that participate in and modulate immune responses (59). Previous and ongoing studies suggest an additional role for *Tlr2* in facilitating epithelial cell barrier integrity and remodeling after a significant immune response has occurred (60–63). As such, we suggest a model where *Tlr2* expression increased at 0.5 dpc in response to the presence of sperm that may modulate epithelial cell integrity, thereafter, inducing remodeling in damaged cells at 1.5 and 2.5 dpc. Future studies need to be conducted to further reinforce this hypothesis.

Further indications of a pro-inflammatory condition induced by sperm at 0.5 dpc followed by epithelial cell remodeling at 1.5 dpc were observed in our luminal proteomics data. We observed an increase in NFκB fluorescent signal at 0.5 dpc, indicating conditions favorable for pro-inflammation. Previous work both *in vivo* and *in vitro* in the uterus and oviduct, respectively, indicate sperm have the capacity to induce immune-related responses (39, 64, 65). In the uterus, observations suggest a hostile, phagocytic environment to remove excessive and dead sperm (64). Our findings suggest an equally hostile response to allogenic sperm in the oviduct at 0.5 dpc. However, this finding is in conflict with prior *in vitro* studies in the bovine oviductal epithelial cell (BOEC) culture model, in which sperm bind and induce anti-inflammatory cytokines, such as *TGFB1* (transforming growth factor β1) and *IL10*, while decreasing pro-inflammatory transcripts such as *TNF*___ (tumor necrosis factor L) and *IL1B* in the BOECs (39, 64). It is possible that this discrepancy could be due to differences between 1) *in vivo* vs. *in vitro* models or 2) murine vs. bovine model organisms. Surprisingly, our luminal proteomics data suggests an exacerbated pro-inflammatory state in the SO condition, inducing greater dysregulation of pro-inflammatory pathways and epithelial cell remodeling.

At 1.5-3.5 dpc, oviductal transcriptional profiles were similar to each other compared to that of 0.5 dpc. During this preimplantation developmental period (1.5-3.5 dpc), embryos transit from the IA to the IU region (51, 66). It indicates that there could be a critical transition of transcripts from 0.5 dpc to other timepoints when the embryos are present at 1.5-3.5 dpc. Therefore, our observations suggest that the oviduct provides an adaptive response in a unique manner during fertilization/preimplantation development, facilitating dynamic selection processes in the presence of gametes and embryos. At 1.5 dpc, 2-cell embryos were in the isthmus region. All embryos at later developmental stages 1.5-2.5 dpc were stalled in the lower isthmus and subsequently the UTJ region between 2.5 and 3.0 dpc. At 3.5 dpc, all embryos have transited from the oviduct to the uterus. Nutrients such as pyruvate, lactate, and amino acids are present in the oviductal fluid in several mammalian species (67–69). After fertilization, zygotes acquire pyruvate and lactate for their energy source (70). Then, the metabolism profile shifts from oxidative to glycolytic metabolism at later stages of preimplantation development (71, 72). Here, we found that upregulated DEGs at 0.5 dpc were enriched for several energy metabolism BPs, including pyruvate metabolic, glucose catabolic process to pyruvate, canonical glycolysis, and glycolytic process through glucose-6-phosphose.

These pathways are subsequently downregulated between 1.5-3.5 dpc. We showed that genes involved in these pathways were unique to the IU region with respect to differential expression analysis. Therefore, it is possible that the IU region is priming the environment to adjust to produce specific energy sources required for early and late embryo metabolism as the embryo switches from utilizing pyruvate to utilizing glucose during successive developmental periods in the oviduct.

Lastly, we found that sperm-induced inflammatory conditions were potentially different than those of chronic inflammatory conditions. The inflammatory responses observed in mice and humans exhibit significant differences based on immune cell involvement, mechanisms, and context. In mice, acute inflammation after sperm exposure could be primarily characterized by the activation of neutrophils, which serve as the first responders to injury or foreign bodies. In contrast, human Fallopian tubes with hydrosalpinx conditions displayed chronic inflammatory conditions predominantly involving macrophages and NK cells, suggesting a more complex and sustained immune response. It is also possible that inflammation in the oviduct differs between mice and humans. Understanding these species-specific variations is crucial for developing effective therapeutic strategies, as findings from murine models may not accurately translate to human inflammatory conditions due to the distinct immune dynamics at play.

In conclusion, we have demonstrated through a comprehensive multi-omics study of the oviduct that the transcriptomic and proteomic landscape of the oviduct at 4 different preimplantation periods was dynamic during natural fertilization, pseudopregnancy, and superovulation using three independent cell/tissue isolation and analytical techniques. Most novel findings from this study suggest that: 1) sperm were likely the key mediators in modulating inflammatory responses in the oviduct, potentially priming the oviduct to become tolerable to the presence of embryos, 2) inflammatory cytokine-mediated signals observed were more robustly amplified by the secretory epithelial cells of the oviduct, 3) the oviduct is an immuno-dynamic organ, alternating between a proinflammatory condition at 0.5 dpc to seemingly prioritizing epithelial barrier integrity/rejuvenation and cellular homeostasis between 1.5-2.5 dpc and 4) the oviduct could provide necessary nutrient enrichment in the luminal fluid at different stages of embryonic development. In addition, an initial stage AI learning predictive model has been used to identify influential transcription factors and correlate predictive protein expressions. This initial AI model has recapitulated a similar progressive prediction of TFs correlating to influenced proteins suggesting similar biological/cellular processes as our empirical *in vivo* multi-omics analysis. Overall, our findings reveal an adaptive oviduct with unique transcriptomic profiles in different oviductal regions, along with dynamic proteomics that may be specialized to influence sperm migration, fertilization, embryo transport, and development. These findings and techniques could facilitate developments to ensure a proper microenvironment for embryo development *in vitro*, assisting in establishing standard protocols at the laboratory, agricultural, and clinical levels.

## MATERIALS AND METHODS

### Animals

All animals were maintained at Washington State University and the University of Missouri and were handled according to Animal Care and Use Committee guidelines using approved protocols 6147, 6151, 38927, and 38961. C57BL/6J mice from Jackson Laboratories (Bar Harbor, ME) were used in this study. In all experiments, adult C57BL/6J female mice between 8-16 weeks were used. Some females were naturally mated with fertile C57BL/6J males. Pseudopregnancy was induced by mating females with vasectomized males. The presence of a copulatory plug the next morning was considered 0.5 days post coitus (dpc) for females mated with fertile males and 0.5 days of pseudopregnancy (dpp) for females mated with vasectomized males.

### Hematoxylin and eosin staining

Oviductal tissues were dissected from 0.5-3.5 dpc/dpp of natural fertilization and pseudopregnancy respectively. Oviducts were placed in cassettes individually and submerged in 10% formalin for 12-16 hrs where they were then placed for storage in 70% ethanol the next day at 4°C. Tissue samples were then paraffin-embedded and sectioned at a 5 µm thickness. Sections were stained with hematoxylin and eosin (H&E) using a standard staining procedure as previously described (3).

### Tissue collection for bulk RNA sequencing

Oviductal tissues were collected and stored in pairs (one pair of oviducts per animal) at 0.5, 1.5, 2.5, and 3.5 dpc/dpp of natural fertilization and pseudopregnancy. For 0.5 dpc/dpp tissue, female mice were placed for mating at 21:00h. For 1.5, 2.5, and 3.5 dpc/dpp, female mice were placed for mating between 5-6 p.m. Oviducts were dissected and kept in 1 mL Leibovitz-15 (L15, Gibco, 41300070, ThermoFisher Scientific, Carlsbad, CA) + 1% fetal bovine serum (FBS, Avantor 97068-091, Radnor Township, PA) media for transportation. Before sectioning the oviduct into two regions (Infundibulum + Ampulla (IA), Isthmus + Uterotubal Junction (IU), oviducts were flushed with L15 + 1% FBS media under a 37°C dissecting microscope (Leica MZ10f, Leica Microsystems, Buffalo Grove, IL). The presence of a minimum of 6 embryos per female was confirmed as a benchmark to represent the average litter size. Additionally, embryos were confirmed to be in the correct developmental stage and location in oviductal tissue samples. Then, oviducts were sectioned into IA and IU regions. We defined the IA region by including the infundibulum and cutting at turn three, from turn four to eleven was considered the IU region, which was stripped of uterine tissue enveloping the colliculus tubaris of the UTJ region (5). Tissue samples were placed in a sterile Eppendorf tube and flash-frozen in liquid N_2_. Samples were stored at –80°C for later RNA extraction. All dissections took place between 10:00-13:00h to decrease sample variation. The average time from cervical dislocation of the mouse to flash-freezing tissues was 15:43 (min:sec). The oviducts were collected at the same time points as their dpc counterparts for DPP samples.

### Bulk RNA isolation, sequencing, and analysis

Both tissue and embryo total RNA were extracted utilizing the RNeasy Micro Kit (Qiagen, Germantown, MD) according to the manufacturer’s instructions. DNA digestion was performed with all samples using a Qiagen RNase-free DNase Set (1500 K units). RNA was then shipped to the University of California San Diego (UCSD) for quality control, library preparation, and sequencing. RNA integrity (RIN) was verified using TapeStation for a minimum RIN value of 7. RNA from this study has an average RIN of 9.04. RNA libraries were prepared using the Illumina Stranded mRNA Prep kit (Illumina Inc., San Diego, CA). Then libraries were sequenced using the Illumina NovaSeqS4 platform with a read depth of 25M reads/sample (*n*=3/region/timepoint), paired-end, and 100bp read length. FASTQ files were then analyzed utilizing BioJupies (73) and an integrated web application for differential gene expression and pathway analysis (iDEP) (74). The quality control, sequence alignment, quantification, differential gene expression (DEG), heatmaps, and pathway and enrichment analyses were performed using default settings as indicated in BioJupies and iDEP web tools (73, 74). In brief, FASTQ files were pseudoaligned, and DEGs were determined using DESeq. DEGs were then plotted as Principal Component Analysis (PCA) and heat maps through BioJupies. In some cases, read counts or reads per kilobase of transcript per million mapped reads (FPKM) were exported from BioJupies and imported into iDEP.92 for further pathway and KEGG analyses. InteractiVenn (95) was used to generate common/overlap gene lists between different regions and timepoints. To validate that our isolation method and RNA-seq data analysis pipeline are reproducible with the previous report (2) we evaluated the gene expression profiles of IA and IU regions from estrus samples (n=3 mice/region). In agreement with the previous findings (2), principal component analysis (PCA) plots showed that the IA and IU regions segregate from each other along the PC1 axis (74.3%) with respect to estrus (data not shown). Similar to the previous report, there was a significant indication of a region-specific expression of large subsets of genes.

### Single-cell isolation, library preparation, and single-cell RNA-sequencing

Another set of mice was used for single-cell isolations and scRNA-seq analysis. Mating and tissue collection protocols were similar to bulk RNA isolation described above, with the exception that female mice were superovulated using the protocol described previously (75) to ensure sufficient numbers of female mice at each time point could be harvested for single cell isolation and library preparation within the same day (n= 3-4 mice/group). Superovulation (SO) was performed by intraperitoneal injection of 5 IU pregnant mare serum gonadotropin (PMSG, Prospect HOR-272, East Brunswick, NJ). Forty-eight hrs after PMSG injection, females were injected with 5 IU of human chorionic gonadotropin (hCG, Prospect HOR-250). Immediately after the hCG injection, females were placed in fertile male cages for mating. Oviducts were collected at 0.5, 1.5, and 2.5 dpc and dissected into IA and IU regions before single-cell isolation. For the control group, oviducts were collected 16 hrs post hCG injection. Trypsin-EDTA (0.25%, MilliporeSigma, T4049) was used for oviductal cell isolation using our previously described method (3). The final cell concentration was targeted for 8,000 cells/run. Cell singlets were captured for the library preparation using 10X Chromium Controller and Chromium Next GEM Single Cell 3’ GEM, Library & Gel Bead Kit v2 (10X Genomics, Pleasanton, CA). Libraries generated were then evaluated for quality using Fragment Analyzer (Agilent, Santa Clara, CA). Libraries were sequenced using Illumina HiSeq4000 at the University of Oregon, targeting 400 million reads/run, paired-end, and 100 bp read length. scRNA-seq web summary output for each dataset is listed in Supplemental Table S2.

### scRNA-seq analysis

Scanpy was used to analyze the scRNA-seq data. The generated loom files were read in as anndata objects and concatenated into a master anndata object. Preprocessing and quality control were performed similarly to the methods described in Scanpys clustering tutorial (76) and Seurat’s clustering tutorial (77). Filtered out were cells expressing fewer than 200 genes, genes expressed in fewer than 3 cells, doublets (cells/droplets with counts for greater than 4,000 genes), and cells with greater than 5% mitochondrial gene counts. Total counts were normalized to 10,000 for every cell, and log transformed. Highly variable genes were then identified using scanpy’s “highly_variable_genes” function with default parameters. Effects of mitochondrial gene expression and total counts were regressed out, and the data was scaled to unit variance and a mean of zero. Dimensionality reduction was first achieved through principal component analysis with Scanpy’s default parameters. To achieve further dimensionality reduction, a neighborhood graph of cells was computed, utilizing the top 40 principal components (PCs) and a neighborhood size of 10, then embedded utilizing Uniform Manifold Approximation and Projection (UMAP), using the default parameters in Scanpy. Clustering of cells was achieved through Leiden clustering at a resolution of 0.1.

Established marker genes were used to identify clusters as specific cell types: pan-epithelial (*Epcam*+), secretory (*Pax8*+), ciliated (*Foxj1*+), leukocytes (*Ptprc*+), antigen-presenting cells (*Cd74+*), monocytes and macrophages (*Ms4a7+ Cd14+*), T-cells (*Cd3d+, Cd3g*), natural killer and NKT cells (*Nkg7+, Klrb1c+*), B-cells (*Cd79a+, Cd79b+*), granulocytes (*S100a8+, S100a9+*), Neutrophils (*Ly6g+*), fibroblasts and stromal (*Pdgfra*+*, Twist2*+*, Dcn*+*, Col1a1*+), and endothelial (*Pecam1*+). Subsets containing only specific cell types (e.g., secretory cells), treatments (e.g., control, 0.5, 1.5, and 2.5 dpc), or regions (e.g., IA and IU) were created for specific downstream analyses and analyzed through the same process as above with identical parameters.

### Oviductal luminal fluid collection for luminal proteomic characterization

Oviducts were collected as pairs at estrus, 0.5, 1.5, and 2.5 dpc/SO of natural fertilization and superovulated fertilization, respectively. The estrus stage was determined by performing a vaginal lavage, followed by H&E staining. Datasets from the natural cycle and SO allowed us to directly compare the impact of exogenous hormone treatments on protein abundance and profile distinct from the physiological levels of hormones. In this context, our SO approach facilitates multi-dimensional analysis comparisons among naturally cycling bulk RNA-seq, SO scRNA-seq, and natural luminal proteomic biological replicates, enhancing confidence between different methods. This experimental design also reflects adaptive responses in the oviduct during natural fertilization and preimplantation development, influenced by PMSG and hCG treatments at both RNA and protein levels. Furthermore, SO is commonly used in female reproduction to synchronize estrus cycles in animals, thus reducing variables at each collection timepoint.

For estrus SO, oviducts were collected the day after hCG injections between 10:00-13:00 h. The presence of cumulus mass cells containing oocytes was also confirmed. For 0.5 dpc/SO tissue, female mice were placed for mating at 9 p.m. For 1.5 and 2.5 dpc/SO, female mice were placed for mating between 5-6 p.m. Oviducts were dissected and washed in a petri dish containing a 25 µL drop of phosphate-buffered saline (PBS) + HALT (1x) (Thermo Scientific, 78440). Once transported to a dissection scope, oviducts were then moved to a fresh adjacent 25 µL drop of PBS + HALT (1x). Inserting a dulled 30G needle and syringe into the UTJ, each oviduct was subsequently flushed with 100 µL PBS + HALT (1x), for a total sample volume of 225 µL. Next, we observed, staged, and removed oocytes/embryos present in the sample drop via mouth pipetting ensuring to take the least amount of sample fluid possible. The presence of a minimum of 6 embryos per female was confirmed as a benchmark to represent the average litter size. Once oocytes/embryos were removed, we placed the sample drop in a 1.5-mL Eppendorf tube and centrifuged at 2200*g* for 15 min to remove any additional cell debris or blood cells that may be present after flushing. The supernatant was removed, and we performed additional centrifugation at 5000*g* for 10 min. Once centrifugation was complete, the supernatant was placed/pooled and flash frozen with liquid N_2_. Pooled samples (n=3 mice/timepoints) were stored at –80°C. Every sample submitted for LC-MS/MS contained 5 paired oviduct flushes at each respective timepoint/condition. The average collection time from cervical dislocation to flushing was 10:47 (min:sec) before subsequent centrifugations. Once all samples were collected, they were shipped on dry ice overnight to Tymora Analytical Operations (West Lafayette, IN) to perform LC-MS/MS analysis.

### ELISA analysis

Enzyme-linked immunoassay (ELISA) was utilized to establish *in vivo* translation of pro-inflammatory cytokine IL1β. Pairs of oviduct tissue from each biological replicate from the IU region were frozen individually, and subsequent protein extraction/tissue disruption of a single IU oviduct at each timepoint/condition was utilized. 2.5 µg total protein concentration from each biological replicate was administered in the assay. Three technical replicates per individual biological replicate at each timepoint/condition were analyzed. Oviductal tissues were collected as pairs (one pair of oviducts per animal) at 0.5 dpc/dpp and 1.5 dpc/dpp of natural fertilization and pseudopregnancy, respectively. Oviducts were dissected and kept in 1 mL Leibovitz-15 (L15, Gibco, 41300070, ThermoFisher Scientific) + 1% fetal bovine serum (FBS, Avantor 97068-091, Radnor Township, PA) media for transportation. Before sectioning the oviduct into two regions (IA and IU), oviducts were flushed with L15 + 1% FBS media under a 37°C dissecting microscope (Leica MZ10f, Leica Microsystems, Buffalo Grove, IL). The presence of a minimum of 6 embryos per female was confirmed as a benchmark to represent the average litter size. Additionally, embryos were confirmed to be in the correct developmental stage and location in oviductal tissue samples. Oviducts were sectioned into IA and IU regions. Tissue samples were stored individually (the pair of oviducts were stored individually) in two separate sterile Eppendorf tubes and flash-frozen in liquid N_2_. Samples were stored at –80°C for later protein extraction (TPER Tissue Protein Extraction Reagent (Thermo Scientific: 78510) + HALT (1x)). IL1β ELISA (ab197742, abcam, Waltham, MA) was performed according to the manufacturer’s protocol.

### NF**κ**B immunofluorescent staining

NFκB immunofluorescent (IF) staining was performed to evaluate the degree of inflammation activation in oviductal cells during fertilization. Following dissection, oviducts were placed in cassettes individually and submerged in 10% formalin for 12-16 hrs where they were then placed for storage in 70% ethanol the next day at 4°C. They were subsequently processed and embedded in paraffin wax. In short, oviductal tissues were sectioned to 5 µm with a microtome. Sectioned samples were placed on Superfrost Plus Slides and baked overnight on a heat plate at 37°C. Slides were processed in xylene, followed by an alcohol series (100%, 95%, 70%). Antigen retrieval was performed with sodium citrate retrieval buffer + 0.05% Tween-20 (pH 6.0) in a pressure cooker at 90°C for 10 min. Slides were rinsed with 1x TBST (0.05% Tween-20) and blocked with 1x TBST + 5% normal goat serum (NGS) cocktail for 60 min at RT before applying the NFκB primary antibody (Cell Signaling, 6956, 1:1000) in 1x TBST cocktail containing 1% bovine serum albumin (BSA) overnight (12-16 hrs) in a 1x TBST humidified chamber at 4°C. Slides were washed the next morning with 1x TBST before the secondary (1:1500) antibody (Jackson Immuno Research, 115-585-146) 1x TBST + 1% NGS cocktail was applied for 1 hour, covered from light, at RT. Slides were rinsed with 1x TBST before ProLong Diamond Antifade Mounting agent with DAPI (Invitrogen, P36962) was applied. The stained sections were subsequently covered with a glass coverslip. Stained slides were placed at 4°C for at least 24 hrs before imaging immunofluorescence using a light microscope (Leica DMi8, Leica Microsystems). To establish relatively quantitative significance, 15 measurements were taken across two stained representative images from 20× objectives using FIJI software, for a total of 30 measurements per timepoint/condition for relative fluorescent strength.

### RNA *in situ* hybridization

To perform *in situ* hybridization, oviductal tissues were dissected as pairs (one pair of oviducts per animal) from 0.5-1.5 dpc/dpp of natural fertilization and pseudopregnancy, respectively. Oviducts were placed in cassettes individually and submerged in 10% formalin for 12-16 hrs where they were then placed for storage in 70% ethanol the next day at 4°C. Tissue samples were then paraffin-embedded and sectioned at a 5 µm thickness, where subsequent staining of target RNAs was performed utilizing ACDbio RNAscope Multiplex Fluorescent Reagent Kit V2 in accordance with ACDbio recommended protocols. RNAscope probes used were as follows: #317521-*Tlr2-*C1, *#*506391-*Ly6g*-C2, and #318651-*Ptprc*-C3. Images were taken using a Leica DMi8 light microscope with a K8 camera (Leica Microsystems). Three technical replicates per individual biological replicate at each timepoint/condition were analyzed utilizing ImageJ (FIJI) color histogram quantification software, followed by GraphPad and 2-way ANOVA statistical analysis comparing the mean of every row (gene target) to every column (timepoint/condition) to establish significance.

### p38 and phosphorylated-p38 immunoblotting

Immunoblotting was used to establish *in vivo* expression and phosphorylation of p38. Pairs of oviduct tissue from each biological replicate from the IU region were frozen individually, and subsequent protein extraction/tissue disruption of a single IU oviduct at each timepoint/condition was utilized. 12 µg total protein concentration from each biological replicate was administered in the assay. Three individual biological replicates at each timepoint/condition were analyzed utilizing FIJI, GraphPad software, and 2-way ANOVA statistical analysis was performed to establish significance. Oviductal tissues were collected as pairs at 0.5 dpc/dpp and 1.5 dpc/dpp of natural fertilization and pseudopregnancy, as described above. TPER + HALT (1x) cocktail (150 uL) was applied to frozen tissue IU samples, followed immediately by homogenization of cells via a tissue disruptor. Tissues were incubated in the TPER + HALT (1x) cocktail for 2 hrs on ice and were vigorously vortexed every 30 min for approximately 10 sec. HALT protease inhibitor was introduced again at the 1-hr incubation for a final concentration of 1X to ensure continuous inhibition of proteases. Homogenized tissue samples were pelleted at 6,000*g* for 5 min at 4°C, with the subsequent removal of the supernatant, which underwent an additional centrifugation treatment. 10 uL of supernatant was aliquoted out of the cell-debris purified supernatant for BCA protein concentration determination. The remaining supernatant (∼140 uL) was flash frozen in liquid N_2_ and stored at –80°C. Protein supernatants were incubated with 4x Laemmli buffer containing β-mercaptoethanol at a final concentration of 12 µg total protein and heated to 95°C for 7 min. Gel electrophoresis was performed with 1x running buffer (25 mM Tris, 192 mM glycine, 0.1% SDS) at 90V constant for 10 min, then after increasing constant voltage to 120V for approximately 1.5 hrs. Polyvinylidene difluoride (PVDF, Immobilon, IPVH00010) transfer membranes were incubated in methanol for 10 min and washed in 1x transfer buffer (25 mM Tris, 192 mM glycine) before transfer of proteins from Tris-glycine SDS-polyacrylamide gels. The transfer occurred on the ice at 90V for approximately 1.5 hrs. Then membranes were subsequently washed (3x, 5 min) with 1x PBS, 0.05% Tween20 (PBST) at RT before being blocked with 5% non-fat milk (ChemCruz, sc-2325) in 1x PBST for 1.5 hrs at RT. Transfer membranes were thereafter treated with a primary p38 (Cell Signaling, 9212S) (1:1000 dilution) or phosphorylated-p38 (Cell Signaling, 4511S) primary antibody (1:1000 dilution) in 1x PBST + 1% BSA overnight at 4°C. Membranes were washed before the secondary goat anti-rabbit antibody (abcam, ab97051) 1x PBST + 1% non-fat milk cocktail was incubated at RT for 1.5 hrs. Membranes were then incubated with Biorad Clarity Western ECL substrate chemiluminescence kit (Biorad, 170-5060) and subsequently imaged utilizing a Biorad Molecular Imager ChemiDoc XRS+.

### Predictive transformer model for predicting proteomics data from transcriptomics data and identifying key transcription factors

The development of an Integrative AI model involves the utilization of a transformer encoder with a single-head self-attention mechanism. The model’s architecture is depicted in Figure 1. Input data comprises bulk RNA sequencing expressions obtained from naturally fertilized oviduct mice at one of various stages such as estrus, 0.5 dpc, 1.5 dpc, and 2.5 dpc, and the output is the abundance of proteins. Preprocessing steps were applied to raw reads from bulk RNA sequencing and raw protein abundance values, which involved removing genes and proteins lacking recorded expression and abundance values across all time points, respectively. Normalization techniques were employed on the data, including Counts Per Million (CPM) normalization (78) for bulk RNA counts to calculate expression values. Furthermore, these values underwent percentile normalization to be confined in the range [0-1], a critical step for machine learning models to manage exploding/vanishing gradients during training (79). Protein abundance values were normalized using log-min-max within each time point sample. A specific threshold of 0.6/0.8 was defined to label proteins as high abundance or low abundance. The bulk RNA-seq expression matrix, encompassing samples from IA and IU regions, along with an additional feature indicating if a gene is a transcription factor, was incorporated and randomly sampled for data augmentation. The transformer model is equipped with a single-layer transformer encoder featuring a single-head self-attention mechanism to predict the abundancy of proteins (abundant or not) from the input RNA-seq data. The attention mechanism directs its focus towards crucial segments of the input, capturing the key genes (e.g., transcription factors) that influence protein abundance. The augmented RNA-seq data from Estrus, 0.5 dpc, and 1.5 dpc, and the corresponding extracted protein abundance labels, were used to train and validate the transformer model. Subsequently, the model’s performance in predicting the abundance of proteins from RNA-seq data was blindly tested using samples from 2.5 dpc not used in the training and validation. Moreover, the attention matrix derived from the trained transformer model was checked against the results of the differential gene expression analysis and the differential protein abundance analysis to identify significant proteins and the key transcription factors that may influence them across different timepoints.

### Enrichr pathway analysis: Reactome (2022) and Gene Ontology (GO Biological Processes 2023) bulk RNA and luminal proteomics analysis

Differentially expressed gene lists were generated for bulk RNA and luminal proteomic data analysis utilizing Biojupies differential expression software and Perseus software. Differentially abundant proteins were transformed with a Gaussian normal assumption before being subjected to two/single-tail *t*-test statistical analysis in Perseus. For a greater description of this integration, refer to the respective methods section below. Differential gene/protein lists were generated with an FDR < 0.05 before subsequent lists were submitted to Enrichr for biological pathway analysis. Reactome (2022) and GO Biological Processes 2023 tables were generated and utilized in combination as each database establishes pathway *p-*value significance differently.

### Gene ontology (GO) scRNA Analysis

Differentially expressed genes were identified using scanpy’s “highly_variable_genes” function with default parameters. Generated DEG sub lists containing up– and downregulated genes with a log_2_FC ≥ 1 or ≤ –1 respectively were then filtered for genes/proteins. The filtered gene/protein lists were submitted to Enrichr for Gene Ontology enrichment analysis. Exported data were plotted utilizing R studio via ggplot (80), InteractiVenn, and Perseus software.

### Gaussian Normal Distribution Assumption (continuous probability distribution)

The Gaussian normal distribution is a mathematical assumption. This mathematical assumption is based on the existence of a continuous random variable. We assume that any single empirically measured protein value (random variable) will not yield the same empirical measurement if subsequent repeated measurements are taken on the same sample. Therefore, we assume that any repeated empirical measurement of any one protein will adhere to a distribution rather than being an absolute measurement.

We applied this assumption to our pooled oviductal luminal protein biological replicates to extend our analysis with respect to utilizing statistical tests to identify significantly altered protein abundances during preimplantation development. However, we limited our assumption range to one standard deviation above and below all empirically measured protein values. Applying these parameters generates two additional artificial but probable values centered around the true empirical measurement. For example, statistical *t*-tests carried out with this integration will assign empirical measurements as the means for statistical comparisons. This transformation allowed for the establishment of significance between proteins with pooled (3 pairs of oviducts, for a total of 6 pooled oviducts per timepoint/condition) biological replicates at each unique timepoint/condition. Two/single-tailed *t*-tests and PCA were generated with Perseus software to establish significant differences. Significant differentially abundant proteins were assigned and visualized with a Venn diagram produced by the interactiVenn webpage, followed by Enrichr Reactome and GO Biological Processes analysis.

### Data availability

Raw data as fastq files were deposited at Gene Expression Omnibus (GSE270654). Bulk-RNA seq, scRNA-seq, and proteomic data are available in the web search format at https://genes.winuthayanon.com/winuthayanon/oviduct_bulkRNA-seq_pregnancy/, https://genesearch.org/winuthayanon/Oviduct_pregnancy/, and at https://genes.winuthayanon.com/winuthayanon/oviduct_proteins/ respectively.

## ACKNOWLEDGEMENT

The authors thank Kalli Stephens for helping maintain the C57BL/6J mouse colony and Gerardo Herrera for initial analysis of scRNA-seq data. This study is supported in part by the Eunice Kennedy Shriver National Institute of Child Health & Human Development, National Institutes of Health award numbers R01HD097087 to W.W., National Science Foundation grants (DBI2308699 and CCF2343612) to J.C., Washington State University (WSU) Office of Research (RA+$10K) Award to R.M.F., and WSU NIH Protein Biotechnology Training Grant (T32GM008336) to D.J.C.

---

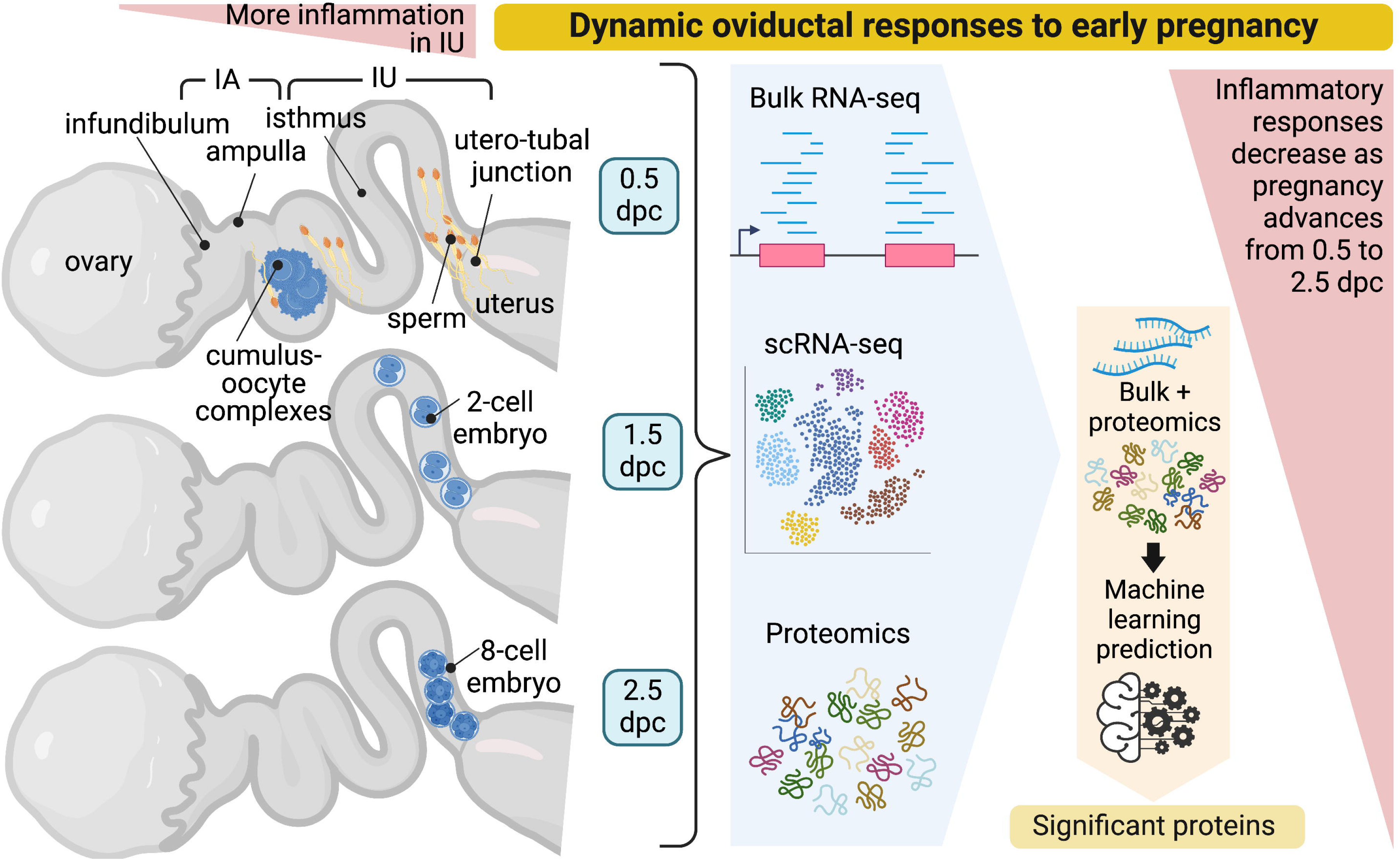

## REFERENCES

1. S. Li, W. Winuthayanon, Oviduct: roles in fertilization and early embryo development. J Endocrinol 232, R1–R26 (2017).

2. E. C. Roberson et al., Spatiotemporal transcriptional dynamics of the cycling mouse oviduct. Dev Biol 476, 240–248 (2021).

3. E. A. McGlade et al., Cell-type specific analysis of physiological action of estrogen in mouse oviducts. FASEB J 35, e21563 (2021).

4. M. J. Ford et al., Oviduct epithelial cells constitute two developmentally distinct lineages that are spatially separated along the distal-proximal axis. Cell Rep 36, 109677 (2021).

5. K. Harwalkar et al., Anatomical and cellular heterogeneity in the mouse oviduct-its potential roles in reproduction and preimplantation development. Biol Reprod 104, 1249–1261 (2021).

6. V. Maillo et al., Oviduct-Embryo Interactions in Cattle: Two-Way Traffic or a One-Way Street? Biol Reprod 92, 144 (2015).

7. C. Alminana et al., Early developing pig embryos mediate their own environment in the maternal tract. PLoS One 7, e33625 (2012).

8. K. F. Lee, Y. Q. Yao, K. L. Kwok, J. S. Xu, W. S. Yeung, Early developing embryos affect the gene expression patterns in the mouse oviduct. Biochem Biophys Res Commun 292, 564–570 (2002).

9. A. Fazeli, N. A. Affara, M. Hubank, W. V. Holt, Sperm-induced modification of the oviductal gene expression profile after natural insemination in mice. Biol Reprod 71, 60–65 (2004).

10. N. Mansouri-Attia et al., Endometrium as an early sensor of in vitro embryo manipulation technologies. Proc Natl Acad Sci U S A 106, 5687–5692 (2009).

11. S. Bauersachs et al., The endometrium responds differently to cloned versus fertilized embryos. Proc Natl Acad Sci U S A 106, 5681–5686 (2009).

12. J. Kropp, H. Khatib, Characterization of microRNA in bovine in vitro culture media associated with embryo quality and development. J Dairy Sci 98, 6552–6563 (2015).

13. S. L. Smith et al., Gene expression profiling of single bovine embryos uncovers significant effects of in vitro maturation, fertilization and culture. Mol Reprod Dev 76, 38–47 (2009).

14. J. L. Tremoleda et al., Effects of in vitro production on horse embryo morphology, cytoskeletal characteristics, and blastocyst capsule formation. Biol Reprod 69, 1895–1906 (2003).

15. H. Niemann, C. Wrenzycki, Alterations of expression of developmentally important genes in preimplantation bovine embryos by in vitro culture conditions: implications for subsequent development. Theriogenology 53, 21–34 (2000).

16. T. P. Fleming et al., The embryo and its future. Biol Reprod 71, 1046–1054 (2004).

17. K. J. Betteridge, D. Mitchell, Direct evidence of retention of unfertilized ova in the oviduct of the mare. J Reprod Fertil 39, 145–148 (1974).

18. D. A. Freeman, G. L. Woods, D. K. Vanderwall, J. A. Weber, Embryo-initiated oviductal transport in mares. J Reprod Fertil 95, 535–538 (1992).

19. G. Lazzari et al., Short-term and long-term effects of embryo culture in the surrogate sheep oviduct versus in vitro culture for different domestic species. Theriogenology 73, 748–757 (2010).

20. B. Leemans et al., Why doesn’t conventional IVF work in the horse? The equine oviduct as a microenvironment for capacitation/fertilization. Reproduction 152, R233–R245 (2016).

21. V. Maillo et al., Spatial differences in gene expression in the bovine oviduct. Reproduction 152, 37–46 (2016).

22. V. Maillo et al., Maternal-embryo interaction in the bovine oviduct: Evidence from in vivo and in vitro studies. Theriogenology 86, 443–450 (2016).

23. W. A. Kues et al., Genome-wide expression profiling reveals distinct clusters of transcriptional regulation during bovine preimplantation development in vivo. Proc Natl Acad Sci U S A 105, 19768–19773 (2008).

24. B. Rodriguez-Alonso et al., Spatial and Pregnancy-Related Changes in the Protein, Amino Acid, and Carbohydrate Composition of Bovine Oviduct Fluid. Int J Mol Sci 21 (2020).

25. B. Rodriguez-Alonso et al., An approach to study the local embryo effect on gene expression in the bovine oviduct epithelium in vivo. Reprod Domest Anim 54, 1516–1523 (2019).

26. K. Smits et al., The Equine Embryo Influences Immune-Related Gene Expression in the Oviduct. Biol Reprod 94, 36 (2016).

27. K. Smits et al., Proteome of equine oviducal fluid: effects of ovulation and pregnancy. Reprod Fertil Dev 29, 1085–1095 (2017).

28. J. A. Weber, D. A. Freeman, D. K. Vanderwall, G. L. Woods, Prostaglandin E2 hastens oviductal transport of equine embryos. Biol Reprod 45, 544–546 (1991).

29. J. A. Weber, D. A. Freeman, D. K. Vanderwall, G. L. Woods, Prostaglandin E2 secretion by oviductal transport-stage equine embryos. Biol Reprod 45, 540–543 (1991).

30. M. A. Marey et al., Local immune system in oviduct physiology and pathophysiology: attack or tolerance? Domest Anim Endocrinol 56 **Suppl**, S204–211 (2016).

31. M. E. Ortiz, P. Bedregal, M. I. Carvajal, H. B. Croxatto, Fertilized and unfertilized ova are transported at different rates by the hamster oviduct. Biol Reprod 34, 777–781 (1986).

32. M. E. Ortiz, C. Llados, H. B. Croxatto, Embryos of different ages transferred to the rat oviduct enter the uterus at different times. Biol Reprod 41, 381–384 (1989).

33. L. A. Velasquez et al., PAF receptor and PAF acetylhydrolase expression in the endosalpinx of the human Fallopian tube: possible role of embryo-derived PAF in the control of embryo transport to the uterus. Hum Reprod 16, 1583–1587 (2001).

34. H. Krzanowska, The passage of abnormal spermatozoa through the uterotubal junction of the mouse. J Reprod Fertil 38, 81–90 (1974).

35. S. Perez-Cerezales et al., The oviduct: from sperm selection to the epigenetic landscape of the embryo. Biol Reprod 98, 262–276 (2018).

36. J. S. Arthur, S. C. Ley, Mitogen-activated protein kinases in innate immunity. Nat Rev Immunol 13, 679–692 (2013).

37. H. Abe, The mammalian oviductal epithelium: regional variations in cytological and functional aspects of the oviductal secretory cells. Histol Histopathol 11, 743–768 (1996).

38. A. S. Georgiou et al., Modulation of the oviductal environment by gametes. J Proteome Res 6, 4656–4666 (2007).

39. M. S. Yousef et al., Sperm Binding to Oviduct Epithelial Cells Enhances TGFB1 and IL10 Expressions in Epithelial Cells as Well as Neutrophils In Vitro: Prostaglandin E2 As a Main Regulator of Anti-Inflammatory Response in the Bovine Oviduct. PLoS One 11, e0162309 (2016).

40. J. Chakraborty, L. Nelson, Fate of surplus sperm in the fallopian tube of the white mouse. Biol Reprod 12, 455–463 (1975).

41. S. Perez-Cerezales, S. Boryshpolets, M. Eisenbach, Behavioral mechanisms of mammalian sperm guidance. Asian J Androl 17, 628–632 (2015).

42. K. Miki, D. E. Clapham, Rheotaxis guides mammalian sperm. Curr Biol 23, 443–452 (2013).

43. R. G. Oliveira, L. Tomasi, R. A. Rovasio, L. C. Giojalas, Increased velocity and induction of chemotactic response in mouse spermatozoa by follicular and oviductal fluids. J Reprod Fertil 115, 23–27 (1999).

44. X. Wang et al., MarsGT: Multi-omics analysis for rare population inference using single-cell graph transformer. Nat Commun 15, 338 (2024).

45. A. Vaswani et al. (2017) Attention is all you need. in *Proceedings of the 31st International Conference on Neural Information Processing Systems* (Curran Associates Inc., Long Beach, California, USA), pp 6000–6010.

46. R. P. Demott, S. S. Suarez, Hyperactivated sperm progress in the mouse oviduct. Biol Reprod 46, 779–785 (1992).

47. Y. Yang et al., Functional roles of p38 mitogen-activated protein kinase in macrophage-mediated inflammatory responses. Mediators Inflamm 2014, 352371 (2014).

48. S. Liu et al., Three Differential Expression Analysis Methods for RNA Sequencing: limma, EdgeR, DESeq2. J Vis Exp 10.3791/62528 (2021).

49. M. Medo, D. M. Aebersold, M. Medova, ProtRank: bypassing the imputation of missing values in differential expression analysis of proteomic data. BMC Bioinformatics 20, 563 (2019).

50. N. D. Ulrich et al., Cellular heterogeneity of human fallopian tubes in normal and hydrosalpinx disease states identified using scRNA-seq. Developmental Cell 57, 914–929.e917 (2022).

51. S. Wang, I. V. Larina, In vivo dynamic 3D imaging of oocytes and embryos in the mouse oviduct. Cell Rep 36, 109382 (2021).

52. L. Ma et al., Association of Epoxide Hydrolase 2 Gene Arg287Gln with the Risk for Primary Hypertension in Chinese. International Journal of Hypertension 2020, 2351547 (2020).

53. S. S. Suarez, The oviductal sperm reservoir in mammals: mechanisms of formation. Biol Reprod 58, 1105–1107 (1998).

54. J. J. Bromfield et al., Maternal tract factors contribute to paternal seminal fluid impact on metabolic phenotype in offspring. Proc Natl Acad Sci U S A 111, 2200–2205 (2014).

55. B. Kaminska, MAPK signalling pathways as molecular targets for anti-inflammatory therapy--from molecular mechanisms to therapeutic benefits. Biochim Biophys Acta 1754, 253–262 (2005).

56. Z. Hu et al., The Repertoire of Serous Ovarian Cancer Non-genetic Heterogeneity Revealed by Single-Cell Sequencing of Normal Fallopian Tube Epithelial Cells. Cancer Cell 37, 226–242.e227 (2020).

57. A. L. Givan et al., Flow Cytometric Analysis of Leukocytes in the Human Female Reproductive Tract: Comparison of Fallopian Tube, Uterus, Cervix, and Vagina. American Journal of Reproductive Immunology 38, 350–359 (1997).

58. W. Xiong, Z. Wang, C. Shen, An update of the regulatory factors of sperm migration from the uterus into the oviduct by genetically manipulated mice. Mol Reprod Dev 86, 935–955 (2019).

59. L. Oliveira-Nascimento, P. Massari, L. M. Wetzler, The Role of TLR2 in Infection and Immunity. Front Immunol 3, 79 (2012).

60. M. T. Abreu, Toll-like receptor signalling in the intestinal epithelium: how bacterial recognition shapes intestinal function. Nat Rev Immunol 10, 131–144 (2010).

61. R. Al-Sadi et al., Bifidobacterium bifidum Enhances the Intestinal Epithelial Tight Junction Barrier and Protects against Intestinal Inflammation by Targeting the Toll-like Receptor-2 Pathway in an NF-kappaB-Independent Manner. Int J Mol Sci 22 (2021).

62. R. Al-Sadi et al., Lactobacillus acidophilus Induces a Strain-specific and Toll-Like Receptor 2-Dependent Enhancement of Intestinal Epithelial Tight Junction Barrier and Protection Against Intestinal Inflammation. Am J Pathol 191, 872–884 (2021).

63. S. Rakoff-Nahoum, J. Paglino, F. Eslami-Varzaneh, S. Edberg, R. Medzhitov, Recognition of commensal microflora by toll-like receptors is required for intestinal homeostasis. Cell 118, 229–241 (2004).

64. M. A. Marey et al., Sensing sperm via maternal immune system: a potential mechanism for controlling microenvironment for fertility in the cow. J Anim Sci 98, S88–S95 (2020).

65. J. E. Schjenken et al., Sperm modulate uterine immune parameters relevant to embryo implantation and reproductive success in mice. Commun Biol 4, 572 (2021).

66. D. Flores, M. Madhavan, S. Wright, R. Arora, Mechanical and signaling mechanisms that guide pre-implantation embryo movement. Development 147 (2020).

67. G. Nieder, C. Corder, Quantitative histochemical measurement of pyruvate and lactate in mouse oviduct during the estrous cycle. Journal of Histochemistry and Cytochemistry 30, 1051–1058 (1982).

68. J. Tay et al., Human tubal fluid: production, nutrient composition and response to adrenergic agents. Human Reproduction 12, 2451–2456 (1997).

69. R. Nichol, R. Hunter, D. Gardner, H. Leese, G. Cooke, Concentrations of energy substrates in oviductal fluid and blood plasma of pigs during the peri-ovulatory period. Journal of Reproduction and Fertility 96, 699–707 (1992).

70. C. Folmes, A. Terzic, Metabolic determinants of embryonic development and stem cell fate. Reproduction Fertility and Development 27, 82–88 (2014).

71. D. K. Gardner, M. Lane, I. Calderon, J. Leeton, Environment of the preimplantation human embryo in vivo: metabolite analysis of oviduct and uterine fluids and metabolism of cumulus cells. Fertil Steril 65, 349–353 (1996).

72. V. Absalon-Medina, W. Butler, R. Gilbert, Preimplantation embryo metabolism and culture systems: experience from domestic animals and clinical implications. Journal of Assisted Reproduction and Genetics 31, 393–409 (2014).

73. D. Torre, A. Lachmann, A. Ma’ayan, BioJupies: Automated Generation of Interactive Notebooks for RNA-Seq Data Analysis in the Cloud. Cell Syst 7, 556–561 e553 (2018).

74. S. X. Ge, E. W. Son, R. Yao, iDEP: an integrated web application for differential expression and pathway analysis of RNA-Seq data. BMC Bioinformatics 19, 534 (2018).

75. W. Winuthayanon et al., Oviductal estrogen receptor alpha signaling prevents protease-mediated embryo death. Elife 4, e10453 (2015).

76. F. A. Wolf, P. Angerer, F. J. Theis, SCANPY: large-scale single-cell gene expression data analysis. Genome Biol 19, 15 (2018).

77. T. Stuart et al., Comprehensive Integration of Single-Cell Data. Cell 177, 1888–1902 e1821 (2019).

78. K. A. Johnson, A. Krishnan, Robust normalization and transformation techniques for constructing gene coexpression networks from RNA-seq data. Genome Biol 23, 1 (2022).

79. L. Huang et al., Normalization Techniques in Training DNNs: Methodology, Analysis and Application. IEEE Trans Pattern Anal Mach Intell 45, 10173–10196 (2023).

80. H. Wickham, Ggplot2: Elegant graphics for data analysis (Springer International Publishing, Switzerland, ed. 2nd, 2016).

