## Supplemental Figure S1 for "Multi-omics analyses and machine learning prediction of oviductal responses in the presence of gametes and embryos"

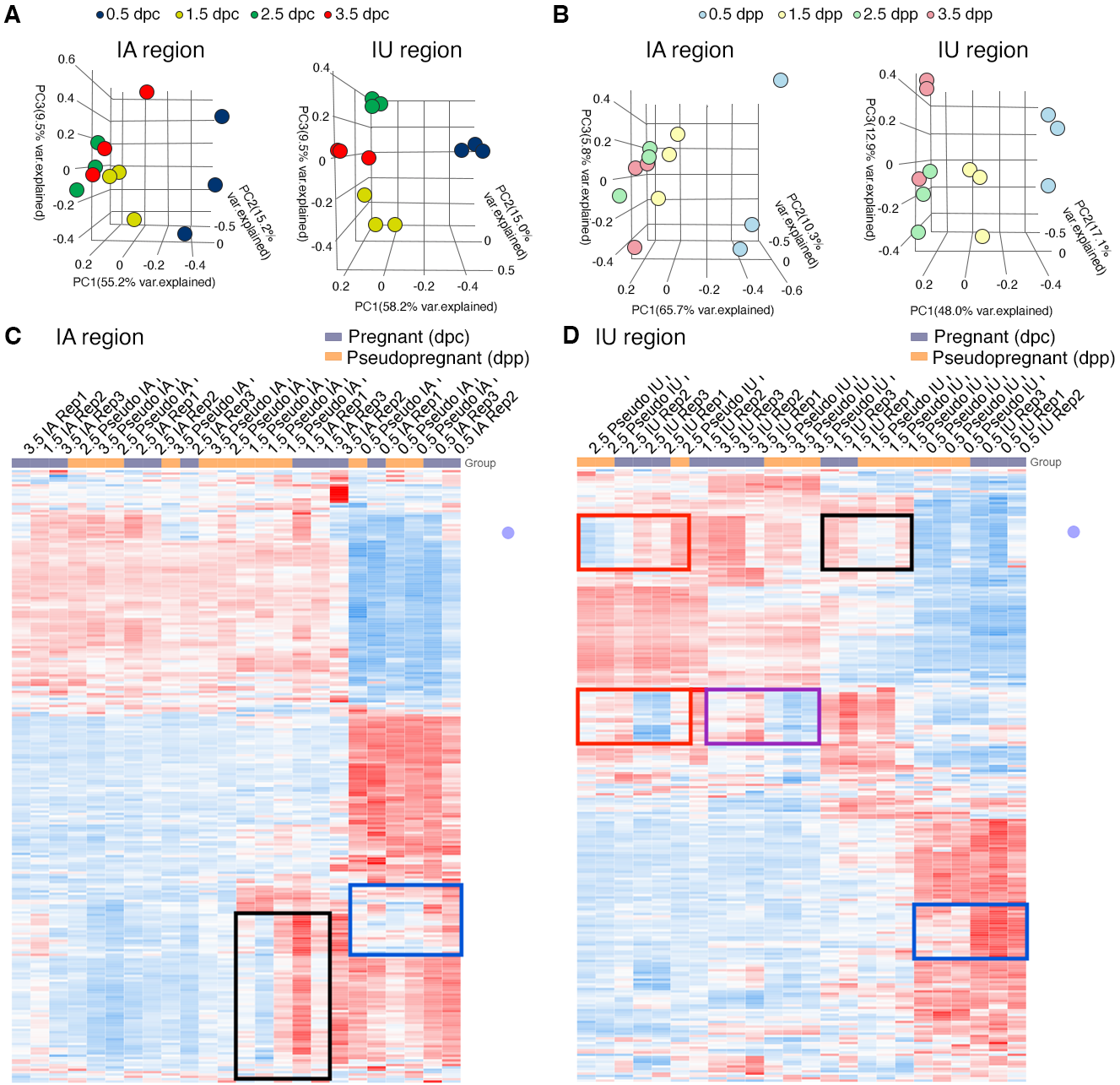

**Supplemental Figure S1.** Principal Component Analysis of top 2500 DEGs identified from bulk-RNA seq of the oviduct collected during **A)** pregnancy at 0.5, 1.5, 2.5, and 3.5 days of pregnancy (dpc) and **B)** pseudopregnancy at 0.5, 1.5, 2.5, and 3.5 days of pseudopregnancy (dpp). Data are separated into infundibulum-ampulla (IA) compared to isthmus-uterotubal junction (IU) regions. **C** and **D)** Heatmap plots of unsupervised hierarchical clustering of top 2500 DEGs in the oviduct during pregnancy (0.5, 1.5, 2.5, and 3.5 dpc) compared to pseudopregnancy (0.5, 1.5, 2.5, and 3.5 dpp) of **C.** IA and **D.** IU regions. Gene expression patterns that differ between 0.5 dpc (blue boxes), 1.5 dpc (black boxes), 2.5 dpc (red boxes), and 3.5 dpc (purple box) compared to the corresponding dpp (n=3 mice/timepoint/region). Blue; downregulated, red; upregulated.
