## Supplemental Figure S2 for "Multi-omics analyses and machine learning prediction of oviductal responses in the presence of gametes and embryos"

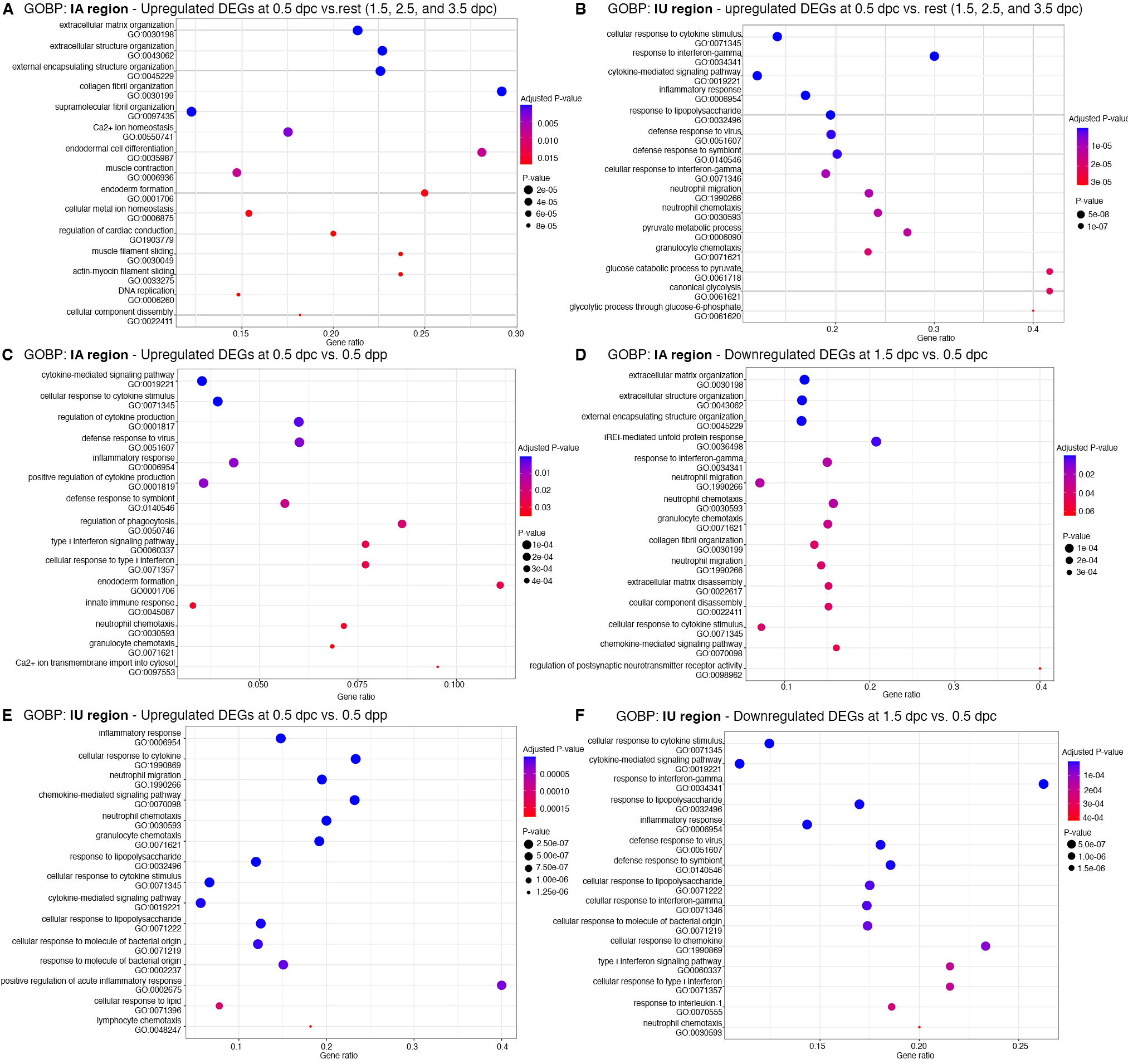

**Supplemental Figure S2.** Dot Plots of biological processes from Gene Ontology (GOBP) analysis from DEGs from bulk RNA-seq analysis at different timepoints. GOBP that were upregulated at 0.5 dpc compared to 1.5-3.5 dpc (0.5 dpc vs. rest) in **A)** IA and **B)** IU regions. GOBP that were **C)** upregulated at 0.5 dpc compared to 0.5 dpp and **D)** downregulated at 1.5 dpc compared to 0.5 dpc in the **C** and **D**) IA region or **E** and **F**) IU region, respectively. GO numbers were indicated within graphs. Gene ratio indicates the ratio of genes that were present in our dataset compared to all genes in the database for each BP.
