## Supplemental Figure S3 for "Multi-omics analyses and machine learning prediction of oviductal responses in the presence of gametes and embryos"

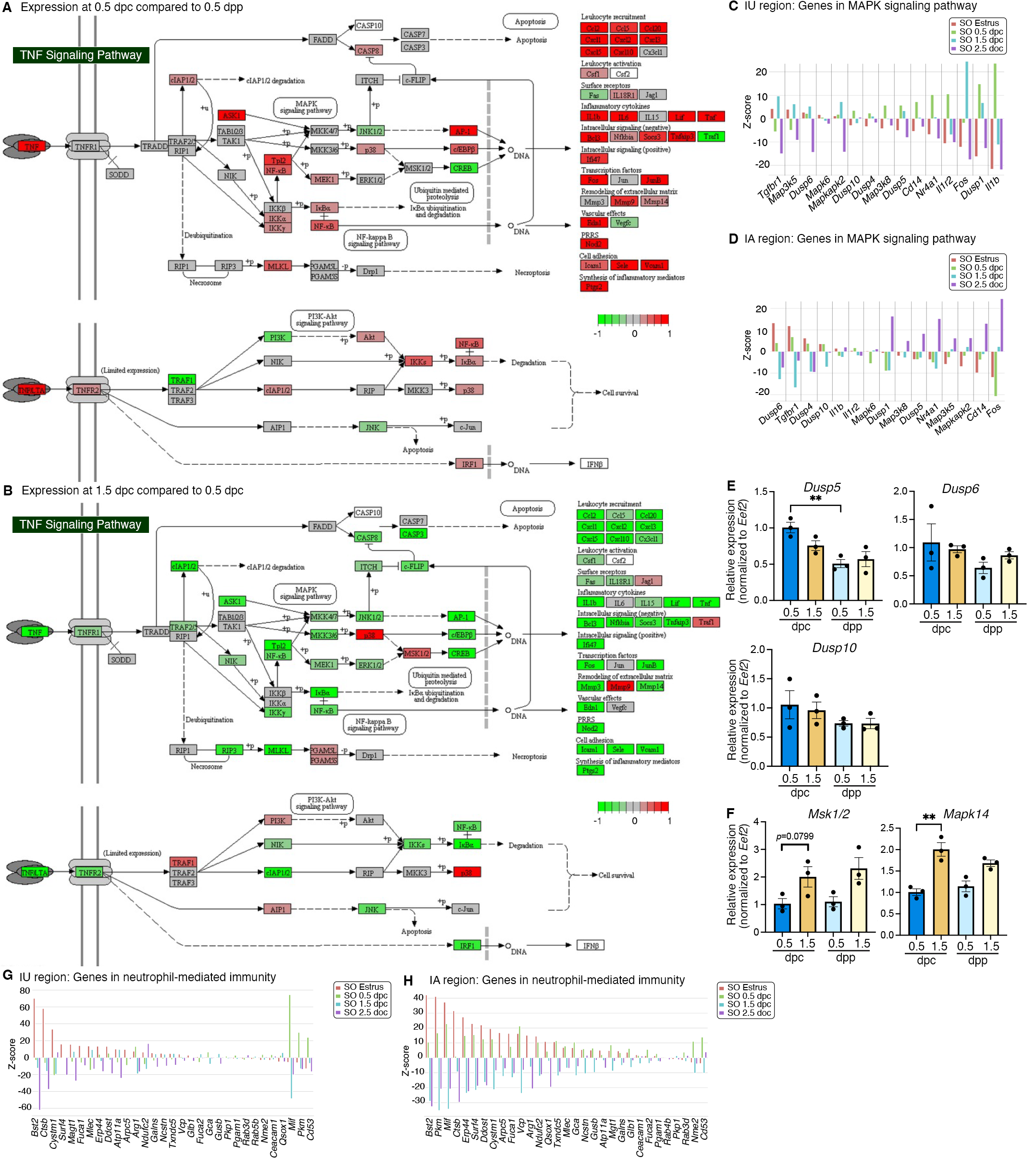

**Supplemental Figure S3.** Analyses of differentially expressed genes that were enriched at different timepoints. KEGG pathway analysis of genes from bulk RNA-seq data in the TGF signaling pathway at **A)** 0.5 dpc compared to 0.5 dpp and at **B)** 1.5 dpc compared to 0.5 dpc. Expression of genes in the MAPK signaling pathways in scRNA-seq dataset in the **C)** IU and **D)** IA region. qPCR validation of **E)** *Dusp5*, *Dusp6*, and *Dusp10* and **F)** *Mak1/2* and *Mapk14* in the whole oviduct at 0.5 dpc, 1.5 dpc, 0.5 dpp, and 1.5 dpp (n=3 mice/timepoint). Expression of genes in the neutrophil-mediated immunity in scRNA-seq dataset in the **G)** IU and **H)** IA region.
