## Supplemental Figure S4 for "Multi-omics analyses and machine learning prediction of oviductal responses in the presence of gametes and embryos"

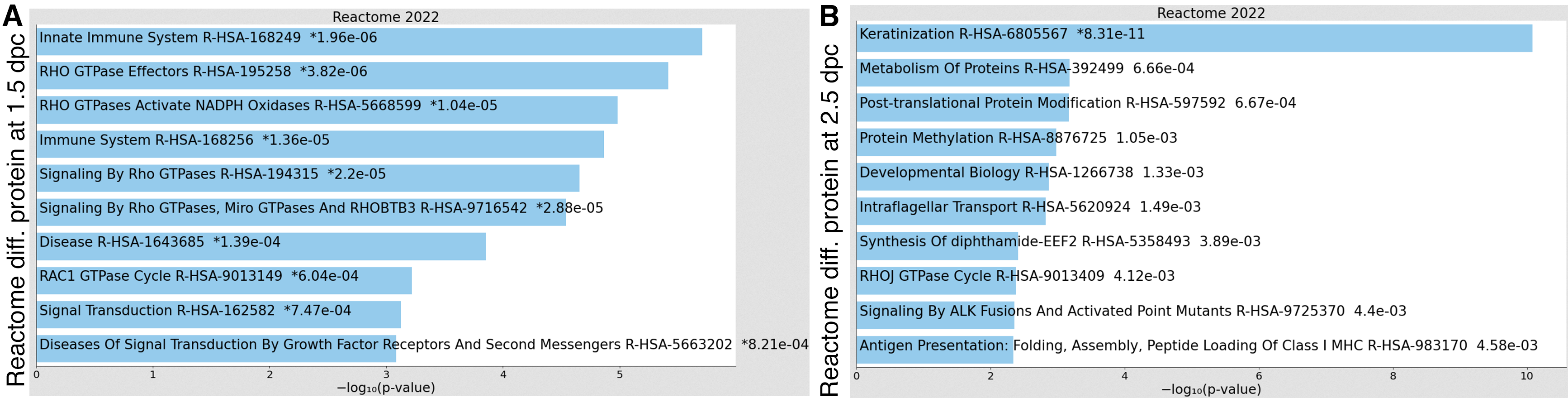

**Supplemental Figure S4.** Enricher Reactome pathway analysis of differentially expressed proteins in the luminal fluid at **A)** 1.5 and **B)** 2.5 dpc.
