## Supplemental Table S1 for "Multi-omics analyses and machine learning prediction of oviductal responses in the presence of gametes and embryos"

**Supplemental Table S1**: The result of predicting protein abundance from bulk RNA-seq data at 2.5 dpc by the transformer model. The model is used to classify proteins into two categories, abundant or not abundant, according to two different thresholds: 0.6 and 0.8, respectively. The model can rather accurately predict the abundant proteins from RNA-seq data.

| Protein Abundance Threshold  (log-min-max normalized) | Accuracy | F1Score | Precision | Recall |
| --- | --- | --- | --- | --- |
| 0.8 | 0.907315 | 0.922326 | 0.985852 | 0.866492 |
| 0.6 | 0.983791 | 0.78453 | 0.972603 | 0.657407 |
