## Supplemental Table S2 for "Multi-omics analyses and machine learning prediction of oviductal responses in the presence of gametes and embryos"

**Supplemental Table S2:** scRNA-seq output for each dataset in this study

| Sample Name | Estimate number of cells | Total reads | Mean reads/cell | Median gene/cell | Median UMI counts/cell | Total detected gene # | Sequencing saturation |
| --- | --- | --- | --- | --- | --- | --- | --- |
| Ctrl_IA | 8,793 | 89M | 10,153 | 1,387 | 2,976 | 19,843 | 32.1% |
| Ctrl_IU | 10,387 | 95M | 9,173 | 967 | 2,194 | 19,285 | 36.6% |
| 0.5_IA | 3,689 | 105M | 28,382 | 1,585 | 3,891 | 19,044 | 64.4% |
| 0.5_IU | 7,551 | 93M | 12,312 | 731 | 1,562 | 18,612 | 67.1% |
| 1.5_IA | 11,760 | 84M | 7,184 | 1,138 | 2,116 | 19,941 | 28.0% |
| 1.5_IU | 10,480 | 84M | 7,989 | 860 | 1,792 | 19,525 | 35.6% |
| 2.5_IA | 4,504 | 107M | 23,870 | 1,616 | 3,590 | 19,250 | 60.5% |
| 2.5_IU | 7,804 | 100M | 12,871 | 928 | 2,178 | 18,350 | 65.0% |
